# Delineation of the Complete Reaction Cycle of a Natural Diels-Alderase

**DOI:** 10.1101/2024.04.18.590041

**Authors:** Laurence Maschio, Catherine R. Back, Jawaher Alnawah, James I. Bowen, Samuel T. Johns, Sbusisiwe Z. Mbatha, Li-Chen Han, Nicholas R. Lees, Katja Zorn, James E. M. Stach, Martin A. Hayes, Marc W. van der Kamp, Christopher R. Pudney, Steven G. Burston, Christine L. Willis, Paul R. Race

**Affiliations:** School of Biochemistry, University Walk, University of Bristol, BS8 1TD, UK; School of Chemistry, Cantock’s Close, University of Bristol, BS8 1TS, UK; Department of Chemistry, King Faisal University, College of Science, Al-Ahsa 31982, Saudi Arabia; Discovery Sciences, BioPharmaceutical R&D, AstraZeneca, Gothenburg, Sweden; School of Natural and Environmental Sciences, Newcastle University, NE1 7RU, UK; Department of Biology and Biochemistry, Claverton Down, University of Bath, BA2 7AY, UK

## Abstract

The Diels-Alder reaction is one of the most effective methods for the synthesis of substituted cyclohexenes. The development of protein catalysts for this reaction remains a major priority, affording new sustainable routes to high value target molecules. Whilst a small number of natural enzymes have been shown capable of catalysing [4+2] cycloadditions, there is a need for significant mechanistic understanding of how these prospective Diels-Alderases promote catalysis to underpin their development as biocatalysts for use in synthesis. Here we present a molecular description of the complete reaction cycle of the bona fide natural Diels-Alderase AbyU, which catalyses formation of the spirotetronate skeleton of the antibiotic abyssomicin C. This description is derived from X-ray crystallographic studies of AbyU in complex with a non-transformable synthetic substrate analogue, together with transient kinetic analyses of the AbyU catalysed reaction and computational reaction simulations. These studies reveal the mechanistic intricacies of this enzyme system and establish a foundation for the informed reengineering of AbyU and related biocatalysts.

## Introduction

The Diels-Alder reaction is a [4+2] cycloaddition between a conjugated diene and a dienophile to form cyclohexenes.^1^ This transformation generates two new carbon-carbon bonds and up to four new stereocentres in a single step, providing an atom-efficient route to the assembly of cyclic and polycyclic products.^2^ Although the Diels-Alder reaction has been extensively studied and widely used in organic synthesis, the existence of naturally evolved protein catalysts for this reaction has until recently remained a matter of contention.^3^ Initially, support for the existence of natural Diels-Alderases was founded exclusively on the observation of putative Diels-Alder reaction products within the scaffolds of molecules isolated from natural sources.^4–6^ More recently, however, studies of prospective Diels-Alderases from natural product biosynthetic pathways have resulted in the formal identification of a number of enzymes shown capable of catalysing [4+2] cycloaddition reactions *in vitro*.^7,8^ These biocatalysts could offer many advantages over synthetic methods, circumventing the requirement for Diels-Alder reactions to be conducted at elevated temperature or pressure, in the presence of additives such as Lewis acids, and with prolonged reaction times, whilst providing more stringent control of product stereochemistry. For these reasons, studies seeking to establish the molecular details of enzyme catalysed Diels-Alder reactions remain an important focus, enabling the development of these biocatalysts as viable tools for ‘green’ synthesis.^9,10^

Naturally evolved [4+2] cyclases are broadly divided into two categories. The first comprises those enzymes that appear to have evolved to catalyse a reaction distinct from cycloaddition, e.g., oxidation, but which serendipitously possess a protein scaffold capable of facilitating cyclisation of the primary reaction product, e.g., solanapyrone synthase.^11,12^The second comprises enzymes which function as explicit [4+2] cyclases,^13–17^ including the Diels-Alderase AbyU.^18^ This biocatalyst generates the spirotetronate skeleton of the antibiotic natural product abyssomicin C (**1**) in the marine actinomycete *Micromonospora maris* AB-18-032 (previously *Verrucosispora maris* AB-18-032). Abyssomicin C readily converts to atrop-abyssomicin C under mildly acidic conditions and the isolation of both natural products has been reported.^19^ The biosynthetic pathway to abyssomicin C is encoded for within a single gene cluster (*aby*), which accounts for ∼60 kb of the *M. maris* genome. This pathway comprises seven polyketide synthase (PKS) modules, distributed across three polypeptide chains (AbyB1 - AbyB3), which function in concert to generate the diketo thiol ester **2** (Figure 1).^20^ This molecule is subsequently liberated from the PKS assembly line via the addition of a glycerate moiety presented on the stand-alone ACP AbyA3, leading to installation of a tetronic acid ring moiety, in a process catalysed by the enzyme AbyA1. Subsequent formation of the exo-methylene (the dienophile) is achieved through acetylation-elimination reactions, catalysed by the AbyA4/AbyA5 enzyme couple to form **3**.^5,21^ Next, AbyU catalyses the key [4+2] cycloaddition reaction between the exo-alkene and the terminal diene, leading to formation of the spirotetronate framework **4** with complete stereocontrol via an *endo* cyclisation. The biosynthesis of abyssomicin C is then completed by selective epoxidation of the 11,12-alkene and an intramolecular attack to form the ether bridge.^22^

**Figure 1.**
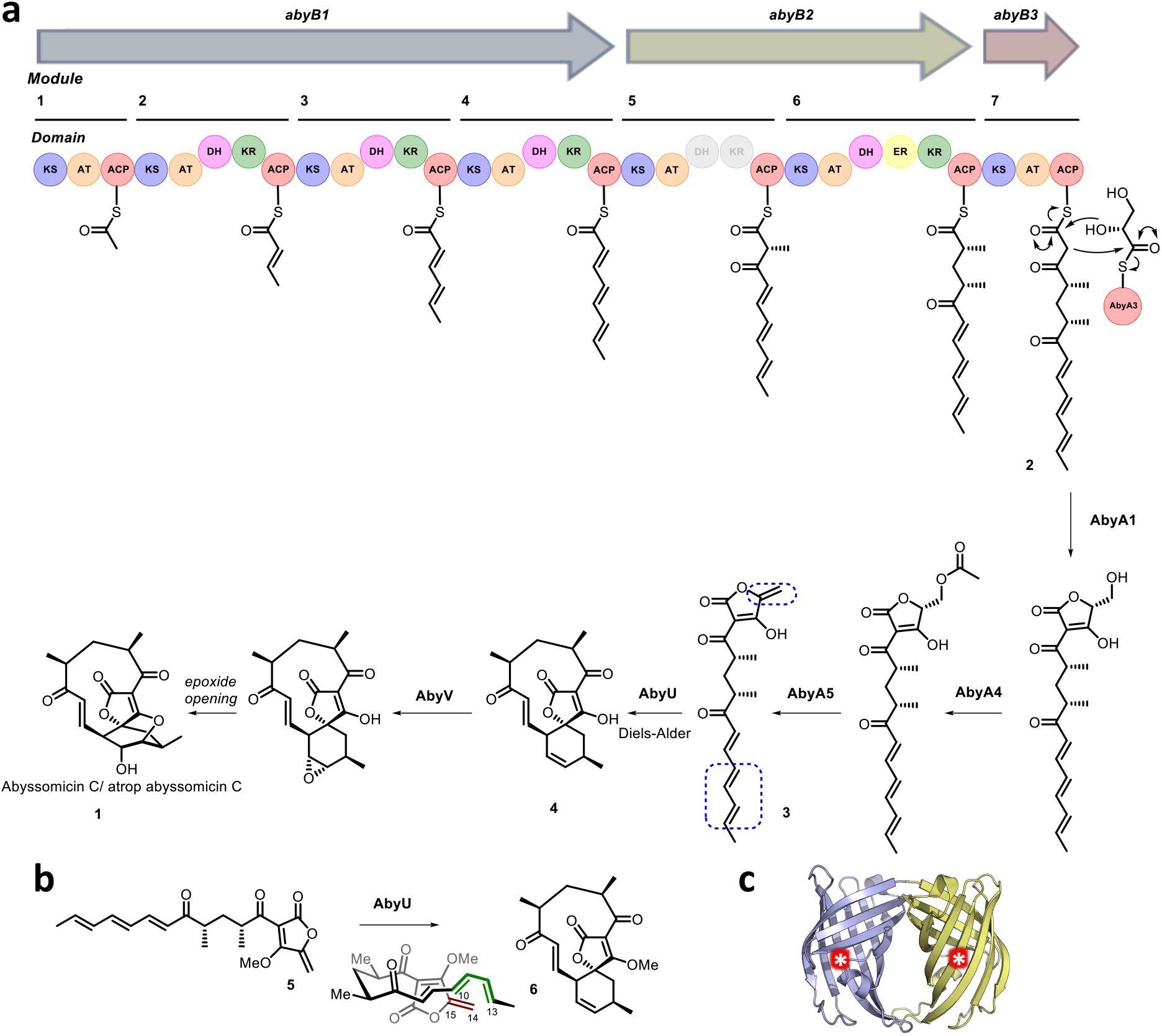
a) Proposed abyssomicin C biosynthetic pathway. The AbyU catalysed [4+2] cycloaddition reaction is highlighted, with the diene and dienophile indicated by blue boxes. b) Chemical structures of the synthesised substrate analogue **5** and (bio)synthesised product **6**. c) X-ray crystal structure of AbyU. Individual β-barrel monomers that comprise the AbyU dimer are coloured blue and yellow, with enzyme active sites indicated by white asterisks in red circles.

Structural studies of AbyU have revealed that this enzyme possesses a distinctive β-barrel fold, wherein the barrel lumen constitutes the active site of the enzyme (Figure 1c). Access to this site is regulated via a flexible capping loop, with catalysis promoted via the conformational constriction of the linear substrate **3**.^18^ However, due to the instability of intermediate **3**, studies have been conducted using the closely related synthetic analogue **5**, with care still required due to competing polymerisation (Figure 1b). This issue, along with similar complications in other experimental systems, has to date precluded the detailed *in vitro* kinetic characterisation of AbyU and related [4+2] cyclases.

Here we resolve these issues and report a molecular description of the complete reaction cycle of AbyU. Our conclusions are based on an X-ray crystal structure of AbyU in complex with a non-transformable substrate analogue, along with transient kinetic studies and *in silico* reaction simulations. Our findings provide important insights into the intricacies of the AbyU catalysed reaction, which is shown to proceed via at least two experimentally verified routes, each arising from a different substrate binding conformation within the enzyme active site. These studies provide new fundamental insights into biocatalytic Diels-Alder reactions and establish a mechanistic framework for the rational reengineering of AbyU and allied biocatalysts.

## Results

### Synthesis of a non-transformable AbyU substrate analogue

To establish the mode of substrate binding in AbyU the substrate analogue **13**, a non-transformable counterpart of **5** was designed for use in co-crystallisation studies. By using analogue **13**, it was hoped that issues of substrate instability which had repeatedly frustrated previous efforts to obtain a structure of the AbyU-substrate complex could be resolved. A resulting co-complex crystal structure would then provide an authentic representation of the AbyU substrate binding mode. The novel enone **13** was prepared from the known lactone **7** (Scheme 1).^18,23^ Phosphonate addition to lactone **7** gave ketone **8**, which, following a Horner-Wadsworth-Emmons (HWE) reaction, generated the required framework for the substrate side-chain as a single diastereomer in 94% yield over the two steps. Dess-Martin periodinane (DMP) oxidation of alcohol **9** gave aldehyde **10** ready for addition of the tetronate and several approaches for the coupling were investigated. The most reliable method proved to be reaction of tetronate **11**^24^ with B-chlorodiisopinocampheylborane [(+)-DIP-Cl] and triethylamine followed by generation of the *exo*-alkene with methyl iodide giving a mixture of diastereomeric alcohols **12**. There was no need to separate the alcohols and the final transformation was oxidation to the required enone **13**, which was fully characterised by spectroscopic methods (SI).

**Scheme 1.**
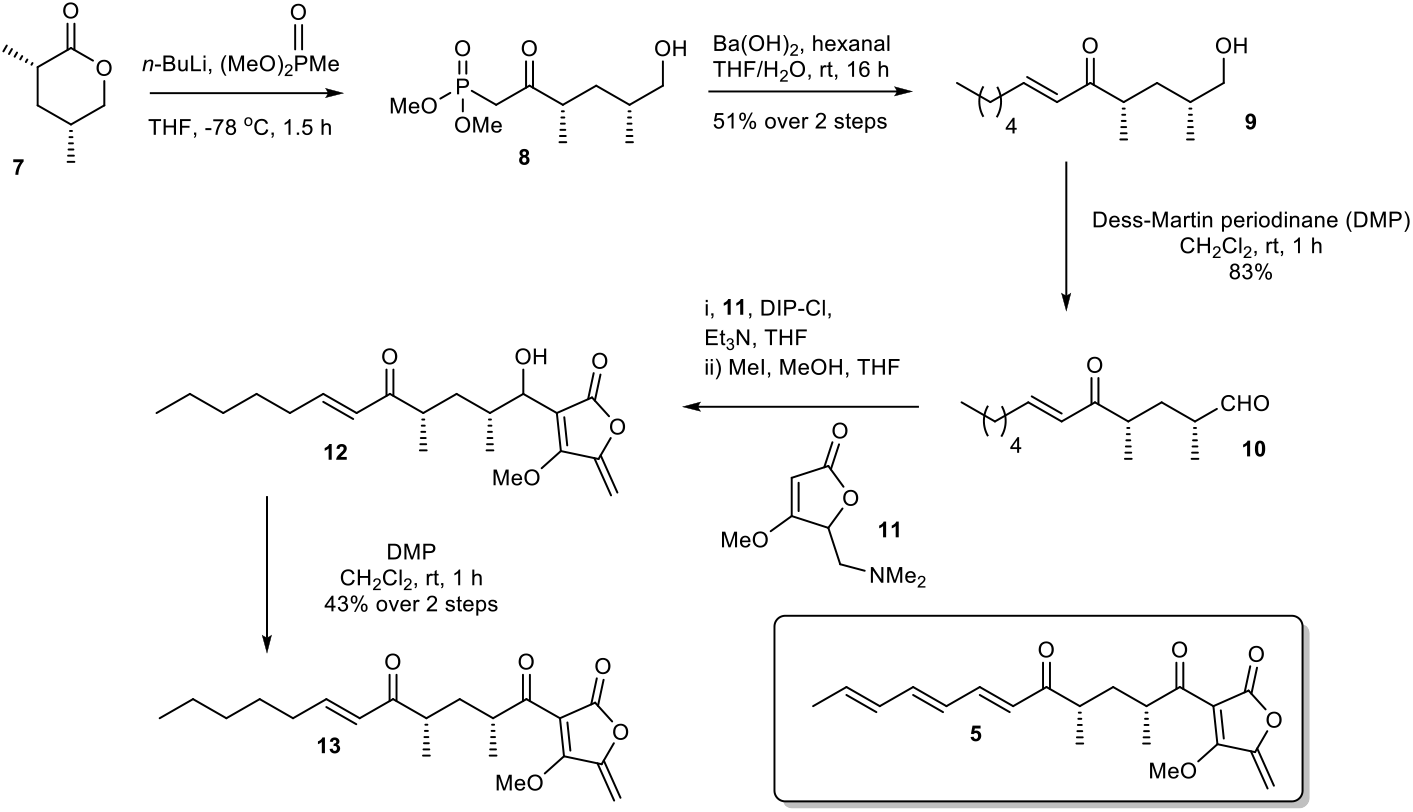
Chemical synthesis of the non-transformable substrate analogue **13**.

### Crystal structure of the AbyU substrate analogue complex

Hexa-histidine tagged AbyU was recombinantly over-expressed in *E. coli* BL21 (DE3) cells and purified to homogeneity as reported previously.^18^ Recombinant AbyU was subsequently subjected to crystallisation screening in the presence of substrate analogue **13**. Resulting crystals underwent X-ray diffraction analysis which yielded a structure of the AbyU-**13** complex to 1.95 Å resolution (Figure 2 and Table S1). Structure determination was performed using molecular replacement employing the published AbyU crystal structure^18^(PDB ID 5DYV) as a search model. Inspection of the AbyU-**13** complex reveals an overall fold comprising a homodimer of two β-barrels consistent with that previously reported for the polypeptide. Both the characteristic hydrophobic active site capping loop and the Glu19-Arg122 salt bridge were unambiguously resolved in the electron density maps. Four copies of AbyU were observed in the asymmetric unit (AU), comprising one intact dimer (between Chains A and C) and two copies (Chains B and D) whose dimer partner resides within a neighboring AU. Electron density corresponding to a single molecule of substrate analogue **13** was observed in the active sites of three of the four copies (Chains A, B and D), with a single HEPES molecule in the fourth (Chain C; Figures 2 and S1-S3). To enable the unambiguous assignment of electron density within the enzyme active site, polder omit maps were generated by excluding local bulk solvent (Figures 2c and S3). The resulting maps show an enhanced density region for each molecule of substrate analogue **13**. The percentage occupancies of **13** differ in each copy of AbyU; 80% in Chain A and 70% in Chains B and D. Superimposition of each of the four copies of AbyU reveals equivalent side chain conformations for all active site residues with the exception of Met126. The side chain of this residue points away from the active site in each of Chains A, B and D, but inwards towards the barrel lumen in Chain C (Figure S2). Given spatial constraints, this reorganisation is undoubtably required to enable substrate binding. In Chain B, electron density is observed consistent with two conformations of the Met126 side chain. Consequently, this residue has been modelled as a dual conformer with differing occupancy levels; 70% for conformation 1, and 30% for conformation 2. In Chains A, B and D a single hydrogen bond of 2.8 Å (oxygen-oxygen distance) is observed between the enzyme and substrate analogue **13**, formed between the lactone carbonyl of **13** and the side chain of Tyr76 (Figure 2b).

**Figure 2.**
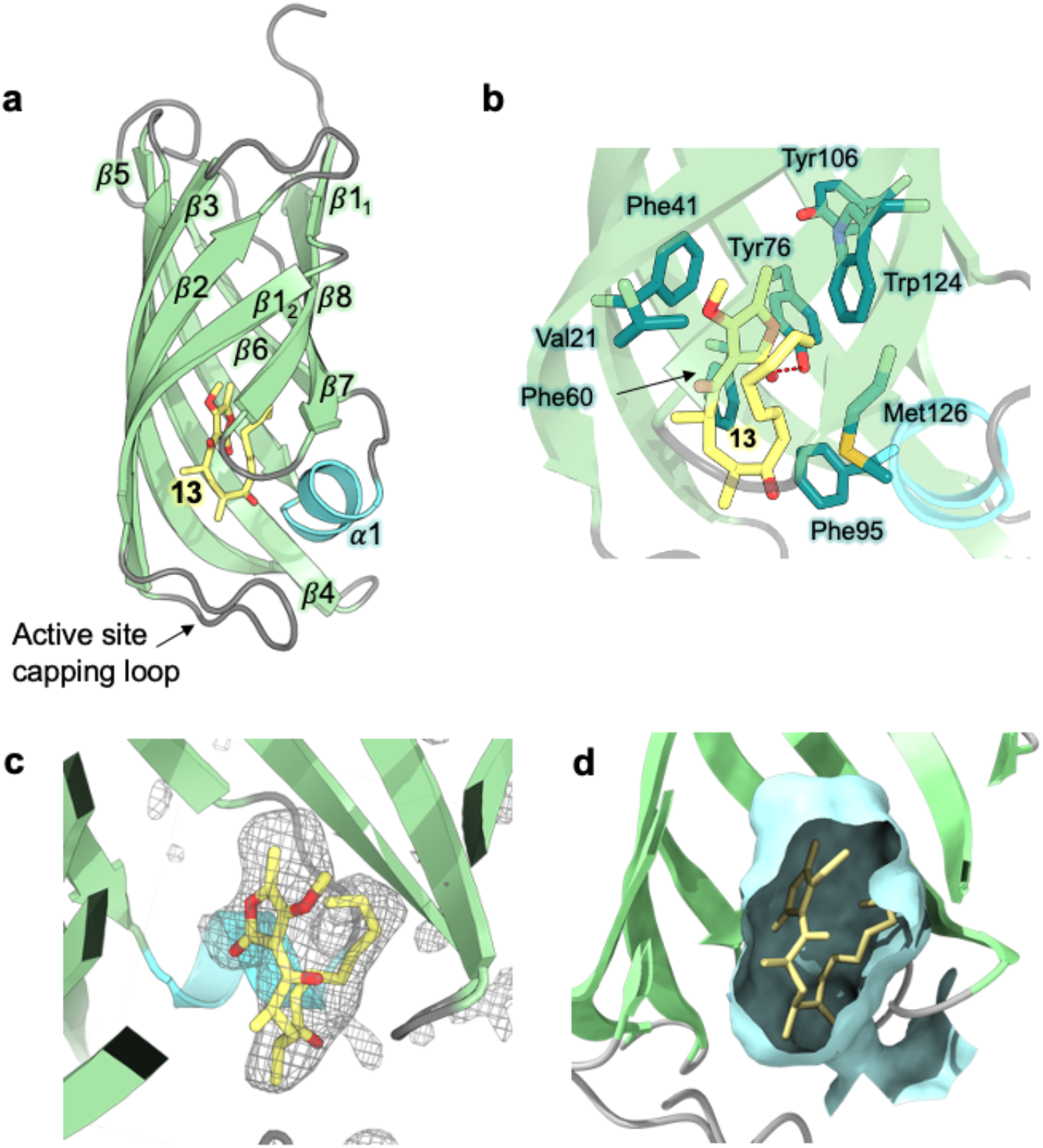
Crystal structure of AbyU in complex with substrate analogue **13**. a) Overall fold of Chain A in ribbon representation (PDB ID 7PXO). Secondary structure elements are coloured as follows: α-helices, turquoise; β-sheets, pale green; loops, grey. The solitary α-helix is labelled 1, with β-sheets labelled sequentially 1-8. The non-transformable substrate analogue **13** is coloured by atom. b) The AbyU active site. The hydrogen bond between Tyr76 and the lactone carbonyl of **13** is indicated by a red dashed line. c) Polder omit map of **13** in the active site of Chain A, contoured at 3 σ. The electron density is shown as a grey mesh. d) Cut through view of the active site showing a surface representation generated using CASTp and coloured pale cyan.

### Transient kinetic studies and observation of multiple substrate binding modes in AbyU

To further investigate the AbyU catalysed reaction stopped-flow kinetic studies were undertaken employing substrate **5**. The absorbance spectrum of **5** reveals a maximum at 325 nm attributed to the triene system of the compound. This feature is lost upon cyclisation and thus constitutes a viable spectrophotometric probe to monitor the AbyU catalysed conversion of substrate **5** to product **6** (Figure 1b). Using an excess of enzyme, analysis of a single turnover of this reaction reveals a highly complex, multi-phasic reaction transient (Figures 3a and S4), from which three distinct exponential phases could be extracted when fitted to Equation 1.

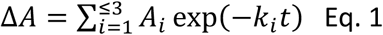

Where *A* is the amplitude, *k* is the observed rate constant (*k*_obs_) for the i^th^ exponential component (up to three) and Δ*A* is the total amplitude change. We assessed the quality of the fit from the plotted residuals. The inclusion of additional exponential phases will inevitably improve the fitting statistics, however, we found that in practice three phases was the minimum required to adequately fit the data.

**Figure 3.**
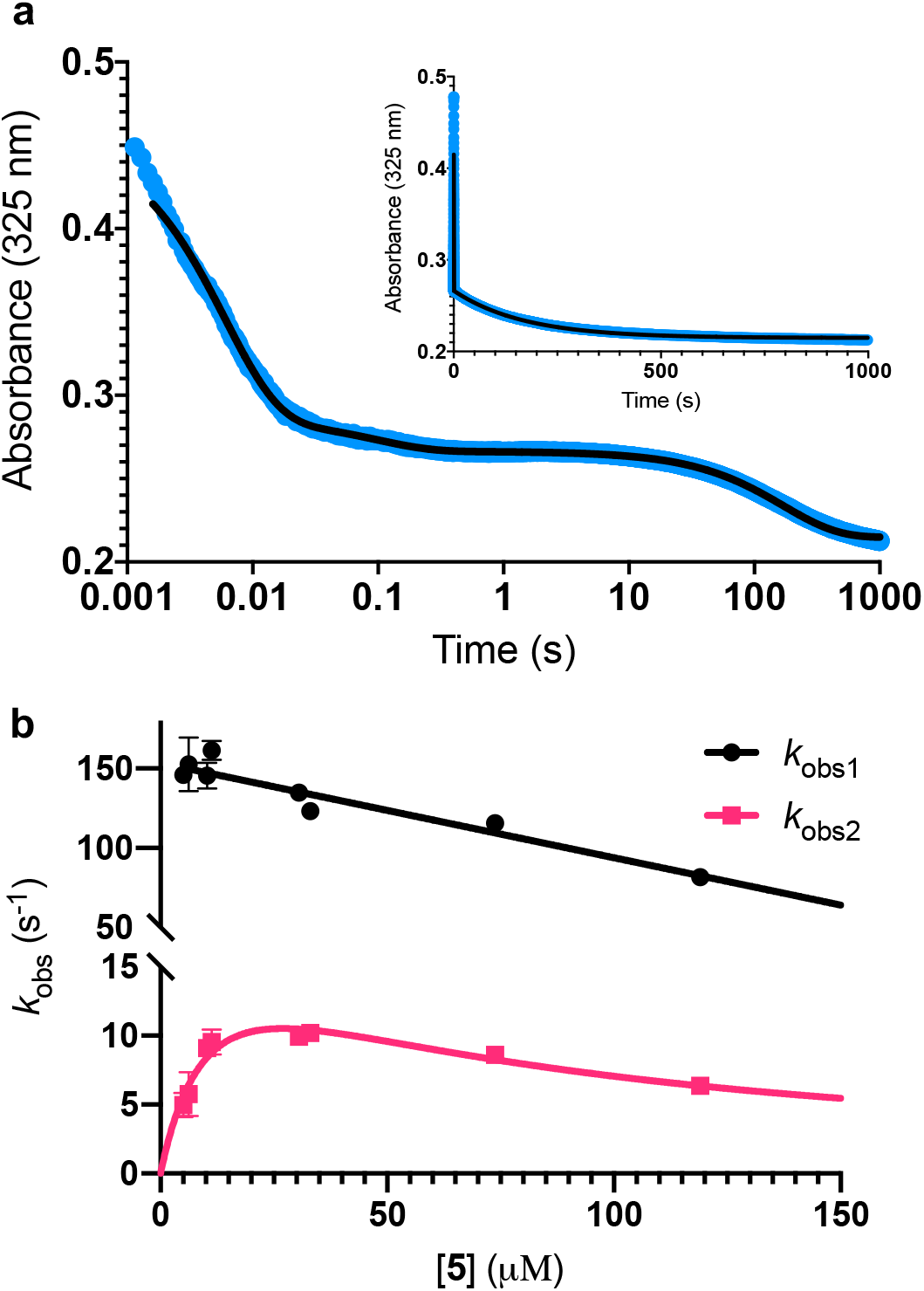
Single turnover analysis of the AbyU catalysed conversion of **5** to **6**. a) Example of the transient observed (blue) in the reaction of 25 μM **5** with 400 μM AbyU, plotted on a logarithmic scale with a linear scale inset for comparison. The transient is fitted to Equation 1 with three exponential terms (black line, see Figure S4 for fitting analysis). b) Concentration dependent analysis of *k*_obs1_ and *k*_obs2_. *k*_obs1_ is fitted to a straight line and *k*_obs2_ is fitted to a substrate inhibition curve (Equation 2).

Multiple exponential phases could indicate the formation of a chemical intermediate, or conformational rearrangement of the enzyme during catalysis.^25,26^ The former can be readily discounted, as [4+2] cycloadditions do not proceed via a sufficiently long-lived intermediate detectable over the timeframe of the experiment.^27,28^ The latter is also improbable, as comparison of our AbyU-**13** complex with that of the unliganded polypeptide reveals the absence of any significant conformational reorganisation during catalysis. Consequently, we suggest that the presence of multiple resolvable exponential phases indicates the existence of multiple possible substrate binding conformations within the AbyU active site. As conformational constriction of the linear substrate is a driver of catalysis, it is conceivable that substrate **5** may be sequestered in different orientations each with a distinct affinity and energetic barrier to catalysis, as has been previously reported in other enzyme systems.^29,30^Given that a single binding mode is observed in our AbyU-**13** crystal structure, we infer that this conformation represents the most energetically favoured configuration.

Analysis of the observed rate constants in each of the fastest two exponential phases, *k*_obs1_and *k*_obs2_, as a function of [**5**] over 10 s (Figures 3b and S5) reveals additional complexity. As the concentration of **5** increases, a concomitant decrease in the rate constants *k*_obs1_ and *k*_obs2_ is observed. This indicates that within each observable phase there exists multiple different binding conformations that react to form product at different rates, i.e., as the concentration of substrate **5** increases, weaker, less productive binding conformations manifest, yielding an apparent reduction in rate. These different conformational populations cannot be individually resolved using ensemble kinetic measurements unless their rate constants differ by at least an order of magnitude, therefore, the observed rate constant reflects average behaviour. The relative equilibrium saturation constants observed for two different binding conformations can be obtained by examining the dependence of *k*_obs2_ on [**5**] (Figure 3b). From Figure 3b, the data present as apparent substrate inhibition. In this case, we can envisage that the ‘slow’ reacting species effectively acts as a weak inhibitor relative to the ‘fast’ reacting species, thus presenting as apparent substrate inhibition. When fitted to a classical substrate inhibition curve (Equation 2), the extracted *K*_m_, 13.2 ± 3.2 μM reflects an approximation of the binding affinity for the faster-reacting conformation(s), whilst the *K*_i_, 54.9 ± 14.5 μM conceptually reflects the binding affinity for slower reacting species.

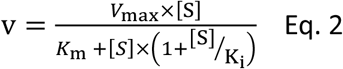

It should be noted that the species reflected by the third exponential phase has a larger equilibrium constant than the faster reacting species, with a rate of reaction approximately 20-fold slower than the steady-state rate of the enzyme (Figure S6). As it represents an insignificant component of the steady-state reaction, we have not analysed this phase any further.

### Modelling and multiscale reaction simulations reveal three key substrate binding orientations

To further investigate the possible alternative binding conformations of substrate **5** in the active site of AbyU, molecular docking, molecular dynamics simulations (with binding affinity calculations) and combined quantum mechanical / molecular mechanical (QM/MM) reaction simulations were performed. As the atropisomer of the cyclised product **6**/**7** is not defined, both possible isomers were investigated, with minimal differences (see Methods for details). Docking of **6**/**7** followed by QM/MM reaction simulations (to form **5**) identified four distinct reactive binding modes of **5** (Figure 4). The reaction simulations indicate that, of these, binding mode A is the most reactive, with cyclisation occurring in an asynchronous concerted manner (Figure S8), consistent with our previous findings.^18^ The reaction barrier predicted for binding mode B is slightly higher than for A.

**Figure 4.**
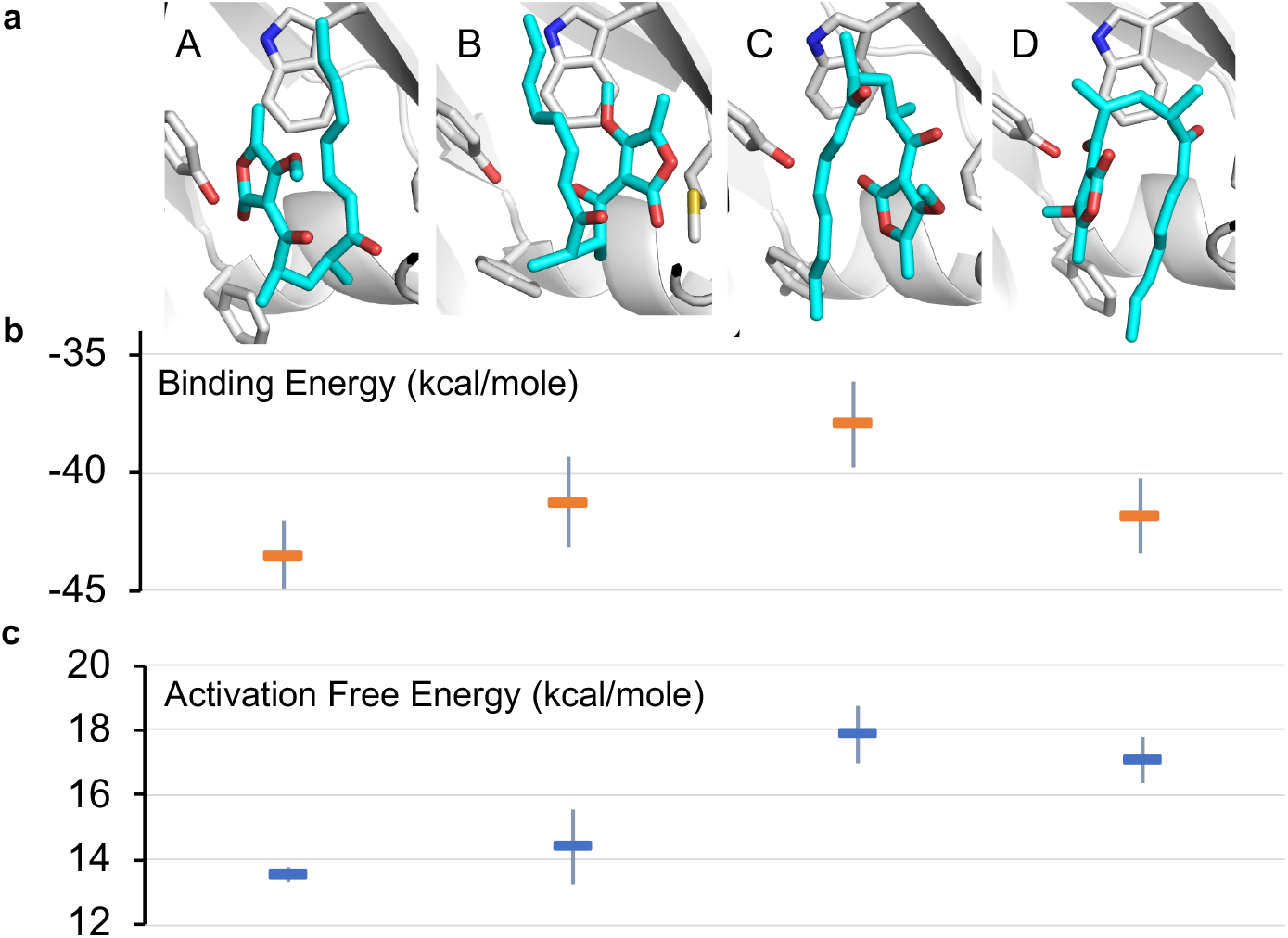
Possible binding modes and respective barriers to catalysis for **5** within the active site of AbyU. a) Representative structures from molecular dynamics (MD) simulations of substrate **5** in each of the four binding modes obtained. b) Average binding energy for each mode (as calculated using the MM-GBSA approach using 1000 snapshots of each mode taken from 10 independent MD runs). c) Predicted Diels-Alder activation free energies for each pose (obtained from 10 independent QM/MM umbrella sampling molecular dynamics simulations using DFTB/ff14SB, with a 7.32 kcal/mole correction for each to M06-2X/6-31+(d,p) level).^18^ Standard deviations are indicated for binding and activation free energies. See Methods for details.

Binding modes C and D both have predicted barriers that are significantly higher than that for mode A and would therefore result in an observably lower rate of turnover. Approximate binding energy calculations (using the MM-GBSA method) indicate that substrate binding mode A is thermodynamically preferred, although modes B and D have comparable binding affinity (Figure 4). This suggests that substrate binding will likely occur in a combination of modes A, B and D, and the substrate may thus be turned over at two distinct rates, consistent with *k*_obs1_and *k*_obs2_, whilst the activation energy barrier for substrate bound in orientation D would be too high for catalysis to occur competitively. *k*_obs3_ therefore represents the rate of dissociation and re-binding of substrate prior to catalysis in a productive conformation.

### Product release is rate limiting in the AbyU catalysed reaction

To formally identify the rate-limiting step in the AbyU catalysed cycloaddition reaction, a multiple turnover kinetics experiment was undertaken. The stopped-flow transients (Figure 5a) reveal two distinguishable burst phases when fitted to Equation 3, a derivative of Equation 1, which incorporates a linear phase to account for multiple turnovers in the steady-state.

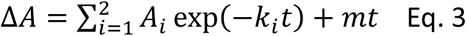

The experimentally observed phases are approximately stoichiometric with the concentration of AbyU (Figures 5a and S6) and therefore are suggestive of product formation via two parallel kinetic routes, as suggested by our single turnover experiments, prior to the rate-limiting step that regulates product release. This is perhaps unsurprising, as the conformationally constricted product must by necessity dissociate from the ‘closed’ active site. Steady-state kinetics reveal the impact that product release has upon the overall rate. The *k*_cat_ of 6.94 ± 0.30 min^−1^ is around 230 times slower than the initial rate of *k*_burst1_, although the *k*_cat_/*K*_m_ of 0.70 ± 0.11 μM^−1^ min^−1^ compares favourably to many other enzymes (Figure 5b). Together, these data demonstrate that product release is rate limiting in the AbyU catalysed reaction.

**Figure 5.**
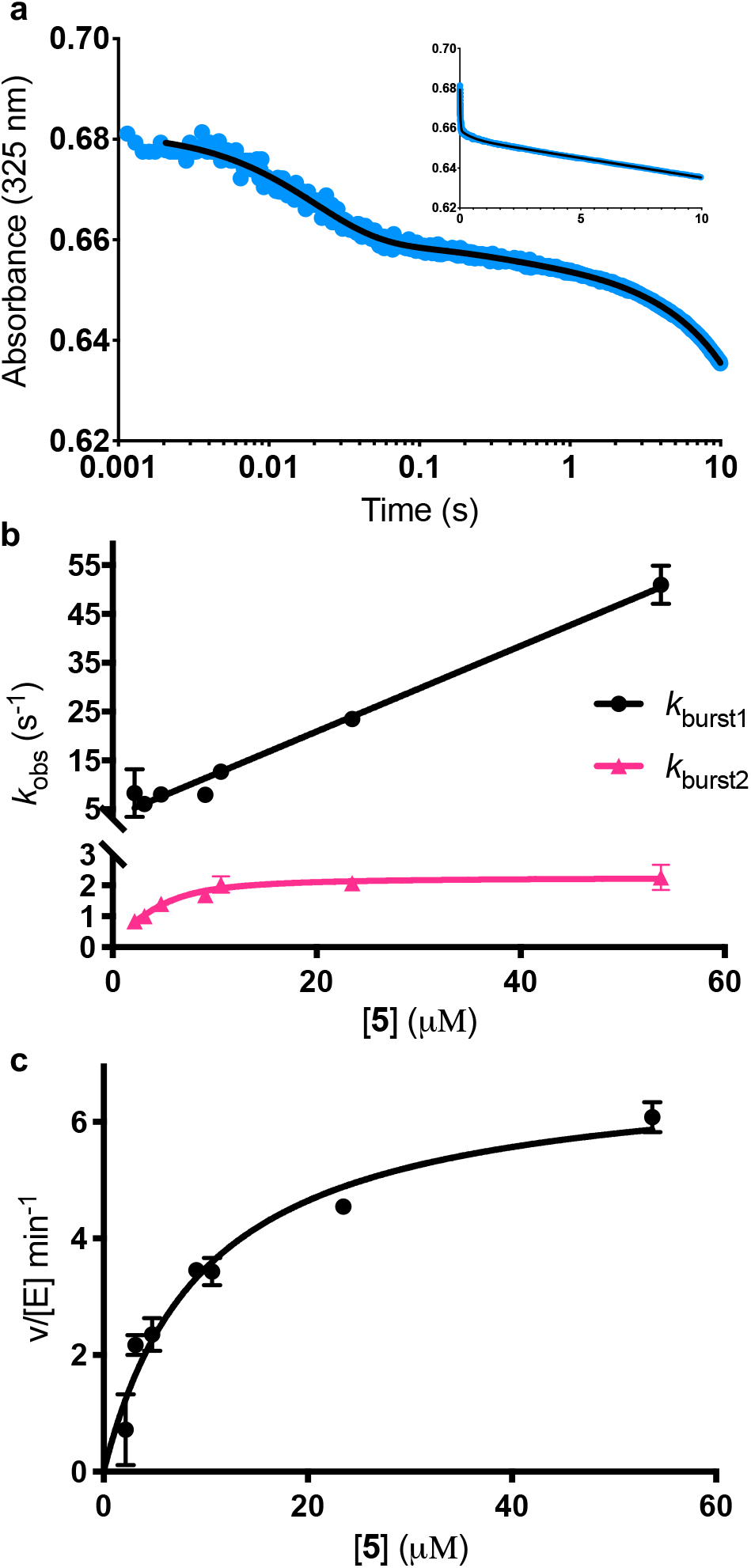
Burst phase characterisation of the AbyU catalysed conversion of **5** to **6**. a) Example of the transient observed (blue) in the reaction of 54 μM **5** with 1 μM AbyU, shown on a logarithmic scale with a linear scale shown inset for comparison. The transient is fitted to Equation 3 (black line, see Figure S9 for fitting analysis). b) The rate of turnover in each burst phase as a function of the concentration of **5**. Data for *k*_burst1_ is fitted to a one-step binding model (Equation 4) and *k*_burst2_ is fitted to a two-step binding model (Equation 5). c) Steady-state kinetics fitted to the Michaelis-Menten equation (Equation S1).

### Cyclisation is driven by substrate rearrangement within the AbyU active site

Further interrogation of the faster of the two phases identified above, *k*_burst1_ (Figure 5c), reveals a linear increase in observed rate constant in relation to the increase in [**5**], and hence a one-step binding mechanism. When the data for *k*_burst1_ are fitted to Equation 4, *k*_on_ and *k*_off_ can be determined to be 0.88 ± 0.03 μM^−1^ s^−1^ and 3.4 ± 0.71 s^−1^, for substrate binding and dissociation respectively, in this conformation.

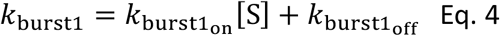

The relationship between the slower burst phase, *k*_burst2_ and substrate concentration, in contrast, fits well to, and is highly representative of, a two-step binding interaction (Equation 5).

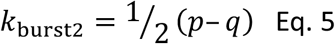

Where:

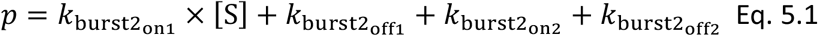

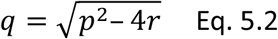

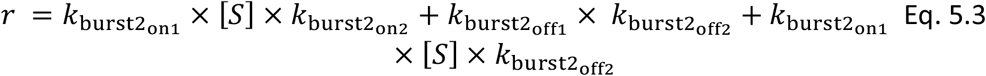

This implies that substrate is initially bound in a sub-optimal conformation, e.g., B in Figure 4, with an on rate of 0.36 ± 0.46 μM^−1^ s^−1^ and an off rate of 0.71 ± 0.62 s^−1^. The substrate then rearranges within the active site to the most productive conformation (A) at a rate of 1.37 ± 2.38 s^−1^prior to catalysis. Although our simulation data demonstrate that bound substrate is unlikely to freely rotate along its horizontal axis (Figure 4), a partial dissociation would be sufficient to facilitate rearrangement, thereby lowering the reaction free energy and promoting catalysis. This process equates to *k*_obs2_ in the single-turnover analysis, and also explains the large disparity between the rate of *k*_obs2_ and *k*_obs3_.

### Complete description of the reaction cycle of AbyU

Deconvolution of our transient kinetic data enables the formulation of a reaction scheme for the AbyU catalytic cycle (Figure 6). These data also permit the derivation of binding constants for each conformation (Table 1). The steady-state rate provides an approximation for product release (*k*_6_), whilst cyclisation occurs rapidly once the substrate is bound (*k*_5_). Our single turnover experiments reveal that each of ES and ES_2_ represent an average of multiple, subtly differentiated substrate binding conformations. These data, together with our burst phase kinetic analyses and *in silico* modelling, reveal that whilst substrate **5** is not selectively bound in a single favoured conformation within the enzyme active site, the internal topology of the AbyU barrel lumen facilitates substrate rearrangement from multiple starting orientations, to a sub-set of reactive binding conformations permitting cyclisation to proceed. The conformation adopted by substrate analogue **13** in our AbyU-**13** crystal structure (Figure 2) is consistent with the substrate binding mode indicated to be most favourable and reactive in our simulations. This mode is thus the most likely optimal binding mode from which catalysis can proceed.

**Table 1.** Experimentally determined rate and binding constants for the cyclisation of **5** by AbyU. *K*_1_ = *k*_-1_ / *k*_1_ and *K*_2_ = *k*_-2_ / *k*_2_. *K*_3_ and K_4_ are calculated in the same way. *K*_app_ = *K*_2_ * *K*_4_.

| Parameter | Constant |
| --- | --- |
| $k_1 (\mu\text{M}^{-1} \text{s}^{-1})$ | $0.88 \pm 0.03$ |
| $k_{-1} (\text{s}^{-1})$ | $3.41 \pm 0.71$ |
| $k_2 (\mu\text{M}^{-1} \text{s}^{-1})$ | $0.36 \pm 0.46$ |
| $k_{-2} (\text{s}^{-1})$ | $0.71 \pm 0.62$ |
| $k_3 (\mu\text{M}^{-1} \text{s}^{-1})$ | $\approx 5.7 \times 10^{-4} \pm 3.8 \times 10^{-5}$ |
| $k_{-3} (\text{s}^{-1})$ | $\approx 6.9 \times 10^{-3} \pm 1.9 \times 10^{-3}$ |
| $k_4 (\text{s}^{-1})$ | $1.37 \pm 2.38$ |
| $k_{-4} (\text{s}^{-1})$ | $0.90 \pm 2.29$ |
| $k_5 (\text{s}^{-1})$ | N.D. (very quick) |
| $k_6 (\text{s}^{-1})$ | $\approx 0.12 \pm 0.05$ |
| $k_{\text{spont}} (\mu\text{M}^{-1} \text{s}^{-1})$ | $1.25 \times 10^{-3} \pm 1.7 \times 10^{-5}$ |
| $K_1 (\mu\text{M})$ | $3.86 \pm 0.94$ |
| $K_2 (\mu\text{M})$ | $1.97 \pm 4.24$ |
| $K_3 (\mu\text{M})$ | $\approx 1.2 \times 10^{-5} \pm 4.1 \times 10^{-6}$ |
| $K_4$ | $0.66 \pm 2.81$ |
| $K_{\text{app}} (\mu\text{M})$ | $1.3 \pm 8.4$ |

**Figure 6.**
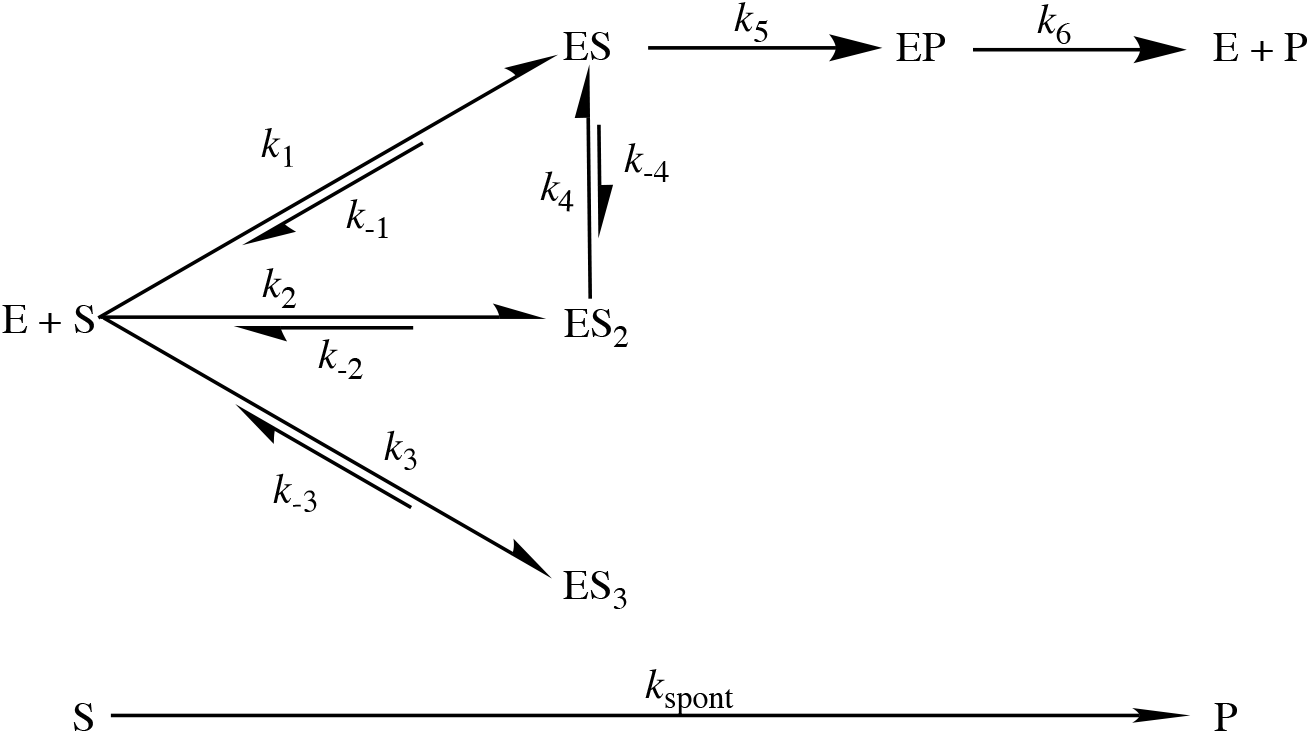
Proposed reaction scheme for the AbyU catalysed Diels-Alder reaction. There are three sets of binding conformations that can be distinguished experimentally, denoted ES, ES_2_ and ES_3_. The intrinsic rate constants calculated for each step are given in Table 1.

## Conclusions

Here we report the delineation of the complete reaction cycle of the natural Diels-Alderase AbyU. Our combined experimental and computational studies reveal a multiplicity of possible binding conformations within the enzyme active site and highlight the occurrence of at least two experimentally validated catalytic routes. AbyU shows specificity for the most productive binding conformations, as exemplified by that observed in our AbyU-**13** complex crystal structure. The architecture of the enzyme active site is sufficiently flexible to enable rearrangement of the substrate without complete dissociation and in the absence of significant conformational rearrangement within the enzyme active site. This permits the *in situ* adoption of energetically favourable reactive binding modes. The rate-limiting step in the AbyU catalytic cycle, product release, is a barrier to optimal catalytic proficiency and presents an opportunity for rational reengineering. Whilst removal of the capping loop in its entirety is unlikely to provide significant benefit, the introduction of mutations in this region that subtly promote loop mobility may enhance the enzyme’s catalytic rate. Attempts to reengineer Diels-Alder biocatalysts should also take into account the multiple binding conformations observed herein and focus on promoting the affinity of substrate binding in the most productive orientations. Together, our findings serve to highlight the mechanistic complexities of biocatalytic Diels-Alder reactions, which incorporate kinetic features that may be readily neglected when applying conventional structure/function experimental techniques.

## Experimental Methods

### Chemical Synthesis

Substrate analogue **5** and Diels-Alder adduct **6** were synthesised as previously described,^18^and as outlined in the SI. Substrate analogue **13** was synthesised as described in the SI.

### Gene Cloning, Expression and Purification of AbyU

AbyU was cloned, over-expressed and purified as previously described,^18^ and as outlined in the SI (Figure S10).

### AbyU Co-crystallisation

For crystallisation experiments AbyU (10 mg/ml) was mixed with substrate analogue **13** (2 mM final concentration, dissolved in 100% acetonitrile) at 20 °C. Conditions supporting the growth of diffraction quality crystals of AbyU were reproduced from Byrne *et al*., 2016^18^ and comprised 0.02 M MgCl_2_, 0.1 M HEPES, 22% Polyacrylic acid sodium salt (MW 5100), pH 7.5. The protein-substrate complex was crystallised using the hanging drop vapour diffusion method, with a 1.5:1:0.5 ratio of protein complex, reservoir solution and seed stock (made from crushing native AbyU crystals, diluted 1:10 with reservoir solution). The resulting crystals were mounted in appropriately sized cryoloops (Hampton Research) and immediately flash-cooled in liquid nitrogen without additional cryoprotection prior to analysis.

### Diffraction Data Collection and Structure Determination

Diffraction data were collected at Diamond Light Source, UK, on beamline I24. Data were auto-processed, merged and scaled with Xia2/3dii^31,32^ using the Diamond Light Source Information System for Protein Crystallography Beamline (ISPyB) managed online environment. All subsequent downstream processing was performed using the CCP4 suite of programs.^33^The structure of the AbyU-**13** complex was initially determined in space group *P*12_1_1 using molecular replacement in MOLREP,^34^ using Chain A of the native AbyU structure (PDB ID 5DYV) as a search model. The structure was determined to 1.95 Å resolution, with four molecules in the asymmetric unit. Three of the molecules of AbyU contained the non-transformable substrate analogue **13** in their active site, the fourth contained a HEPES molecule. Two additional HEPES molecules were identified in the asymmetric unit. Iterative rounds of manual model building and refinement using COOT^35^and Refmac5^36^ were used to refine the structure. Data collection, phasing, and refinement statistics for the AbyU-substrate complex are provided in Table S1. Omit maps were created to improve visualisation of the substrate electron density using the Polder Maps program^37^ as implemented by the Python-based Hierarchical Environment for Integrated Xtallography (Phenix) suite of programs.^38^ Protein structure graphics were prepared using PyMOL Molecular Graphics System, Version 2.5.2. The structure of the AbyU-**13** complex has been deposited in the PDB with the accession code 7PXO.

### Stopped-Flow Experiments

Stopped-flow (SF) experiments were performed using a Hi-Tech Scientific KinetAsyst Stopped-Flow System spectrophotometer (TgK Scientific, Bradford on Avon, UK). Both syringe 1 and 2 were washed with ddH_2_O and SF buffer (18 mM Tris-HCl, 135 mM NaCl, 10 % acetonitrile, pH 7.5). 10 % acetonitrile was maintained in both syringes at all times to ensure solubility of the substrate. The stop-flow was zeroed and references were taken using SF buffer in each syringe for burst phase kinetics. Purified recombinant AbyU in SEC buffer (20 mM Tris-HCl, 150 mM NaCl, pH 7.5), was diluted to a final concentration of 800 μM AbyU in SF buffer. This solution of AbyU was placed in syringe 1 to obtain references for the single turnover experiments. For the multiple turnover experiments, AbyU was added to syringe 1 at a concentration of 2 μM (diluted in SF buffer). Likewise, substrate **5** was diluted in SF buffer to double the reaction concentration and placed in syringe 2. 50 μL from each syringe was injected into the mixing chamber and measurement immediately commenced. Reactions were observed using a logarithmic timebase over either 1000 s or 10 s. For the 1000 s experiments, the timeframe was split into 15 cycles, each with 200 points, each of which was distributed in a logarithmic manner. This ensured a high number of readings in the first seconds of the experiment. It did not prove tractable to measure each reaction over 1000 s as over this timeframe **5** can undergo significant background cyclisation to form **6**, precluding accurate repeats at a consistent starting concentration. This was compounded by the weak affinity of substrate **5** for the plastic components in the SF system, rendering it impossible to precisely replicate the concentration of **5** in the reaction chamber over multiple syringes. To measure the uncatalysed rate of substrate cyclisation, the stop flow was zeroed and references were taken using SF buffer in both syringes. Varying concentrations of substrate **5** were added to syringe 2 and measurements were taken using a linear time base over 10 seconds. All reactions were performed at 22 °C.

### Data Fitting and Analysis of the Stopped-Flow Transients

Results were fitted using Graphpad Prism. Data obtained before 2 ms was discarded, reflecting the dead-time of the instrument. Data were fit to equations as outlined in the text. The 10 s single turnover kinetic data were fitted to two exponentials with a sloping baseline (Equation 3). This fit did not, however, reliably determine the angle of the slope observed, due to the shallow nature of the slope and bias introduced by the exponential phases. To circumvent this, the data between 3 s and 10 s were separately fitted to a straight line to determine the initial rate of *k*_3_. In order to fully analyse the change in the reaction rate with respect to the concentration of **5**, 3-6 transients at each substrate concentration were taken and the rate and amplitude parameters derived from the fit appropriate for each transient (Figure S4 and S9). The background rate of reaction (Figure S11) was removed from the steady-state reaction and from the linear portion of *k*_3_. The other exponential phases occur too rapidly for the background rate of reaction to have an appreciable influence on the data. The amplitude of the change in absorbance was converted to concentration using the following measured absorbance coefficient for substrate **5**: ε325 nm = 13200 M^−1^cm^−1^ and the following measured absorbance coefficient for product **6**: ε325 nm = 1085 M^−1^ cm^−1^. The rate and amplitude parameters were averaged across all transients from the same concentration of **5**, before plotting against the initial concentration of substrate. Initial concentrations of substrate **5** were calculated from the absorbance at t = 0, to account for uncatalysed substrate turnover, and the tendency of the substrate to have a weak affinity with the plastic present in the stop-flow apparatus. Once plotted the data were subsequently fitted as is outlined in the text. In order to calculate the rate of spontaneous cyclisation by **5**, the transients were each fitted to a straight line. The slope of the line was determined for each substrate concentration and averaged over 3 – 5 repeats.

### Molecular Docking for Binding Mode Investigation

Molecular docking was used to obtain different binding poses for the possible reaction products of the AbyU substrate analogue **5** (Figure S7). Starting conformations (in PDB format) of the two product atropisomers **6** and **7** were generated in Chem3D, optimised using the built-in MM2 minimiser. The coordinates for the enzyme were taken from chain A of the crystal structure of AbyU (PDB ID: 5DYV),^18^ removing the buffer molecule HEPES bound in the active site. AutoDockTools 1.5.6 was used to create the input files for each protein/product docking by assigning AutoDock atom types to the relevant atoms as well as defining rotatable bonds. Polar hydrogens are required in the input structures of both the ligand and protein to order to determine hydrogen bond donor/acceptors. Since the protein crystal structure doesn’t contain hydrogen atoms, polar hydrogens were first added in AutoDockTools, with all residues in their standard protonation states. During docking, the side-chains of residues 76, 95, 124 and 126 (which line the cavity) were treated flexibly; all formally single bonds were defined as rotatable. The standard active torsions detected by AutoDockTools were assigned for the products. Docking was then performed with AutoDock Vina^39^ using a search grid of size 21.4 x 16.5 x 17.6 Å centred on the active site, and an exhaustiveness of 16. Four poses (corresponding to binding modes A-D in Figure 4) for the atropisomer with the keto groups pointing in ‘opposite’ directions and three poses for the alternative atropisomer (**6** and **7**, respectively; a binding mode corresponding to B for atropisomer **7** was not found) were selected for further simulation to predict the reactivity and binding affinity of the different modes. These binding modes are analogous to those found in our previous work,^18^ the difference being that this work now uses the substrate analogue rather than the natural substrate.

### Structural Preparation, Optimisation and Molecular Dynamics of Docked Poses

The Enlighten (www.github.com/marcvanderkamp/enlighten)40 protocol PREP was used to prepare the structures obtained from docking for simulation. PREP first takes the combined protein and docked product structure; it then adds hydrogens (to the protein only) via the AmberTools^41^ program reduce. This assigned the only histidine, His88, as being doubly protonated. Asp43 was protonated, since the predicted p*K*_a_ from PropKa 3.1 was 9.7.^42^ Then, in addition to the crystallographically determined water molecules, a 20 Å solvent sphere was added around the active site (with the AmberTools program tleap), missing heavy atoms (and their hydrogens) were added, and finally the topology and co-ordinate files were generated, describing the resulting system for simulation. The protein was parameterised by the ff14SB force field,^43^ and TIP3P was used as the water model. AM1-BCC partial charges and General Amber Force Field^44^ parameters were generated for each atropisomer using Antechamber. For each of the poses, structural optimisation and Molecular Dynamics (MD) simulation was performed to obtain starting structures for QM/MM umbrella sampling. This was done using the Enlighten protocols STRUCT and DYNAM, using Sander from AmberTools16, with the entire structure treated MM. The STRUCT protocol performs brief simulated annealing and minimisation to optimise the structure. DYNAM then takes the output from STRUCT and performs brief heating to 300 K, followed by MD simulation. Ten repeat DYNAM runs were performed to obtain ten independent 10 ps MD trajectories (2 fs timestep), the final snapshots of which were then used to start umbrella sampling runs. Throughout, all atoms outside of the 20 Å solvent sphere were kept fixed.

### QM/MM umbrella sampling

The reaction involves the concerted formation of two new carbon-carbon bonds, one between C10 and C15 and the other between C13 and C14 (see Figure S7). As simulations are started from the product, the reaction is sampled in reverse, by forcing these two bonds to break. To do this, distances between the respective carbons are gradually increased. A single reaction co-ordinate can be used to describe this, as a weighted average of the two distances:

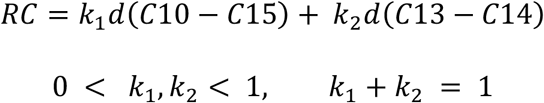

The underlying energy surface, together with the weights, determines the path taken in the reaction simulations, since these determine the relationship that the two distances must satisfy for a given value of the reaction co-ordinate. To ensure that the reaction coordinate leads to sampling along the minimum free energy surface, we first obtained the underlying potential energy surface for the reaction (Figure S8a; Supplementary Methods).

To determine the optimal reaction co-ordinate, umbrella sampling was performed for the atropisomer with both keto groups pointing in the same direction (**6** in Figure S7) in mode A, testing out different weight combinations. Ten repeat umbrella sampling runs were performed for each weight combination, starting from the MD snapshots of the product complex. Umbrella sampling was done at the SCC-DFTB/ff14SB level with the QM region consisting of just the ligand and using 26 windows from reaction coordinate values 1.3 Å to 3.8 Å (i.e. steps of 0.1 Å). A restraint of 200 kcal mol^−1^ Å^−2^and 2 ps of simulation was used for each umbrella sampling window. The simulations were run with the Amber16 program Sander using a 1 fs timestep and keeping all atoms outside the 20 Å solvent sphere fixed. Reaction coordinate values were recorded every 1 fs and used as input for the Weighted Histogram Analysis Method (WHAM)^45^ to obtain the potential of mean force (free energy profile) along the reaction coordinate. In addition to this, the bond distances themselves were recorded every fs to assess sampling quality. Comparing the distances sampled using the 1D co-ordinate to the potential energy surface, the optimal weight combination was found to be *k*_1_=0.3, *k*_2_=0.7, as this ensured that the system sampled the correct (i.e. minimum energy) reaction path through the transition state (Figure S8b). The umbrella sampling protocol described above was then run for the remaining product poses using the optimised reaction co-ordinate, and in each case similar sampling was obtained. For each pose, individual free energy profiles were obtained with WHAM for each of the repeat umbrella sampling runs. The barrier was then calculated for each profile as the difference between the transition state and substrate minimum and the average (and standard deviation) taken from the ten repeats.

### Binding affinity calculations

To calculate an approximate binding affinity for the reactive poses of the substrate that correspond to the different product poses, the MM-GBSA^46^ approach was used. First, an ensemble of enzyme-substrate structures was generated for each pose. For this purpose, 10 starting structures were obtained for each pose by taking them from the final window of the umbrella sampling runs for the corresponding product pose. For each starting structure, after a brief QM/MM optimisation, the Enlighten protocols PREP and STRUCT were run as before and DYNAM was then used to run 100 ps of MD. Structures were saved every 1 ps, so each of the 10 starting structures provided 100 snapshots for an overall ensemble of 1000 structures for each pose. These structures were used for calculating the binding affinity with MM-GBSA using the single trajectory approach. The average and standard deviation of the binding energy over all the snapshots were then reported for each pose.

Representative structures (as shown in Figure 4) were generated for each pose by using cpptraj^47^ to cluster all 1000 structures based on the all-atom RMSD of the substrate, without fitting (using the default hierarchical agglomerative clustering algorithm) into a single cluster and taking the cluster centroid. Note: In Figure 4, results are only reported for one of the atropisomers of each of the binding modes A-D. The trends in reaction barriers were the same for both atropisomers. For modes A and C, the substrate always ended up in a reactive substrate conformation with both keto groups approximately pointing in the same direction at the end of umbrella sampling, regardless of which product it came from; reaction barriers and binding affinities are therefore reported for simulations starting from the corresponding atropisomer, **7**. For mode D, although distinct substrate conformations were obtained for the two atropisomers, the average barrier and binding energy obtained was the same (within 0.2 kcal/mole); only the result obtained when starting from the atropisomer **7** is therefore reported. Finally, for mode B, results are only shown for the alternative atropisomer, since this docking mode was only obtained for that atropisomer.

## Acknowledgments

This work was supported by BBSRC and EPSRC through the BrisSynBio Synthetic Biology Research Centre (BB/L01386X/1), BBSRC grants BB/T001968/1, BB/M012107/1 and BB/M025624/1, through the award of PhD studentships to LM and STJ (EPSRC Centre for Doctoral training in Synthetic Biology, EP/L016494/1 and DSTL) and JIB, SZM and NRL (EPSRC Bristol Centre for Doctoral Training in Chemical Synthesis, EP/L015366/1 and GSK), a BBSRC David Phillips Fellowship to MWvdK (BB/M026280/1), and by AstraZeneca. Computer simulations were conducted using the computational facilities of the Advanced Computing Research Centre, University of Bristol. We thank the Deanship of Scientific Research at King Faisal University, Saudi Arabia, for financial support for JA under Nasher Track (Grant No. 216117). We also thank staff at Diamond Light Source, beamline I24, for assistance with X-ray diffraction data collection, and Rob Barringer for assistance with PDB structure deposition.

## References

1 O. Diels and K. Alder, Justus Liebigs Ann Chem, 1928, 460, 98–122.

2 K. C. Nicolaou, S. A. Snyder, T. Montagnon and G. Vassilikogiannakis, Angewandte Chemie International Edition, 2002, 41, 1668–1698.

3 K. Klas, S. Tsukamoto, D. H. Sherman and R. M. Williams, J Org Chem, 2015, 80, 11672–11685.

4 A. Karine, S. Andrew, K. Jonathan, D. J. Witter, J. P. Van den Heever, C. R. Hutchinson and J. C. Vederas, J Am Chem Soc, 2000, 122, 11519–11520.

5 E. M. Gottardi, J. M. Krawczyk, H. von Suchodoletz, S. Schadt, A. Mühlenweg, G. C. Uguru, S. Pelzer, H. Fiedler, M. J. Bibb, J. E. M. Stach and R. D. Süssmuth, ChemBioChem, 2011, 12, 1401–1410.

6 A. Minami and H. Oikawa, J Antibiot (Tokyo), 2016, 69, 500–506.

7 B. Jeon, S.-A. Wang, M. W. Ruszczycky and H. Liu, Chem Rev, 2017, 117, 5367–5388.

8 K. Zorn, C. R. Back, R. Barringer, V. Chadimová, M. Manzo-Ruiz, S. Z. Mbatha, J. Mobarec, S. E. Williams, M. W. van der Kamp, P. R. Race, C. L. Willis and M. A. Hayes, ChemBioChem, 2023, 24, e202300382.

9 J. Funel and S. Abele, Angewandte Chemie International Edition, 2013, 52, 3822–3863.

10 A. E. Settle, L. Berstis, N. A. Rorrer, Y. Roman-Leshkóv, G. T. Beckham, R. M. Richards and D. R. Vardon, Green Chemistry, 2017, 19, 3468–3492.

11 H. Oikawa, K. Katayama, Y. Suzuki and A. Ichihara, J. Chem. Soc., Chem. Commun., 1995, 1321–1322.

12 J. M. Serafimov, D. Gillingham, S. Kuster and D. Hilvert, J Am Chem Soc, 2008, 130, 7798–7799.

13 T. Hashimoto, J. Hashimoto, K. Teruya, T. Hirano, K. Shin-ya, H. Ikeda, H. Liu, M. Nishiyama and T. Kuzuyama, J Am Chem Soc, 2015, 137, 572–575.

14 Z. Tian, P. Sun, Y. Yan, Z. Wu, Q. Zheng, S. Zhou, H. Zhang, F. Yu, X. Jia, D. Chen, A. Mándi, T. Kurtán and W. Liu, Nat Chem Biol, 2015, 11, 259–265.

15 L. Gao, C. Su, X. Du, R. Wang, S. Chen, Y. Zhou, C. Liu, X. Liu, R. Tian, L. Zhang, K. Xie, S. Chen, Q. Guo, L. Guo, Y. Hano, M. Shimazaki, A. Minami, H. Oikawa, N. Huang, K. N. Houk, L. Huang, J. Dai and X. Lei, Nat Chem, 2020, 12, 620–628.

16 H. J. Kim, M. W. Ruszczycky, S. Choi, Y. Liu and H. Liu, Nature, 2011, 473, 109–112.

17 Q. Zheng, Y. Guo, L. Yang, Z. Zhao, Z. Wu, H. Zhang, J. Liu, X. Cheng, J. Wu, H. Yang, H. Jiang, L. Pan and W. Liu, Cell Chem Biol, 2016, 23, 352–360.

18 M. J. Byrne, N. R. Lees, L.-C. Han, M. W. van der Kamp, A. J. Mulholland, J. E. M. Stach, C. L. Willis and P. R. Race, J Am Chem Soc, 2016, 138, 6095–6098.

19 K. C. Nicolaou and S. T. Harrison, J Am Chem Soc, 2007, 129, 429–440.

20 L. Maschio, A. E. Parnell, N. R. Lees, C. L. Willis, C. Schaffitzel, J. E. M. Stach and P. R. Race, 2019, 617, 63–82.

21 N. R. Lees, L. Han, M. J. Byrne, J. A. Davies, A. E. Parnell, P. E. J. Moreland, J. E. M. Stach, M. W. van der Kamp, C. L. Willis and P. R. Race, Angewandte Chemie International Edition, 2019, 58, 2305–2309.

22 A. J. Devine, A. E. Parnell, C. R. Back, N. R. Lees, S. T. Johns, A. Z. Zulkepli, R. Barringer, K. Zorn, J. E. M. Stach, M. P. Crump, M. A. Hayes, M. W. van der Kamp, P. R. Race and C. L. Willis, Angewandte Chemie International Edition, 62, e202213053.

23 M. B. Andrus, W. Li and R. F. Keyes, J Org Chem, 1997, 62, 5542–5549.

24 L. Montgomery and G. Challis, Synlett, 2008, 2008, 2164–2168.

25 S. Ferrer, I. Tuñón, S. Marti, V. Moliner, M. Garcia-Viloca, À. González-Lafont and J. M. Lluch, J Am Chem Soc, 2006, 128, 16851–16863.

26 J. A. Pavon, B. Eser, M. T. Huynh and P. F. Fitzpatrick, Biochemistry, 2010, 49, 7563–7571.

27 U. Rivero, M. Meuwly and S. Willitsch, Chem Phys Lett, 2017, 683, 598–605.

28 D. Yepes, O. Donoso-Tauda, P. Pérez, J. S. Murray, P. Politzer and P. Jaque, Physical Chemistry Chemical Physics, 2013, 15, 7311.

29 B. Birdsall, J. Feeney, S. J. B. Tendler, S. J. Hammond and G. C. K. Roberts, Biochemistry, 1989, 28, 2297–2305.

30 C. R. Pudney, S. Hay, J. Pang, C. Costello, D. Leys, M. J. Sutcliffe and N. S. Scrutton, J Am Chem Soc, 2007, 129, 13949–13956.

31 G. Winter, J Appl Crystallogr, 2010, 43, 186–190.

32 G. Winter, D. G. Waterman, J. M. Parkhurst, A. S. Brewster, R. J. Gildea, M. Gerstel, L. Fuentes-Montero, M. Vollmar, T. Michels-Clark, I. D. Young, N. K. Sauter and G. Evans, Acta Crystallogr D Struct Biol, 2018, 74, 85–97.

33 M. D. Winn, C. C. Ballard, K. D. Cowtan, E. J. Dodson, P. Emsley, P. R. Evans, R. M. Keegan, E. B. Krissinel, A. G. W. Leslie, A. McCoy, S. J. McNicholas, G. N. Murshudov, N. S. Pannu, E. A. Potterton, H. R. Powell, R. J. Read, A. Vagin and K. S. Wilson, Acta Crystallogr D Biol Crystallogr, 2011, 67, 235–242.

34 A. Vagin and A. Teplyakov, J Appl Crystallogr, 1997, 30, 1022–1025.

35 P. Emsley, B. Lohkamp, W. G. Scott and K. Cowtan, Acta Crystallogr D Biol Crystallogr, 2010, 66, 486–501.

36 G. N. Murshudov, P. Skubák, A. A. Lebedev, N. S. Pannu, R. A. Steiner, R. A. Nicholls, M. D. Winn, F. Long and A. A. Vagin, Acta Crystallogr D Biol Crystallogr, 2011, 67, 355–367.

37 D. Liebschner, P. V. Afonine, N. W. Moriarty, B. K. Poon, O. V. Sobolev, T. C. Terwilliger and P. D. Adams, Acta Crystallogr D Struct Biol, 2017, 73, 148–157.

38 P. D. Adams, P. V. Afonine, G. Bunkóczi, V. B. Chen, I. W. Davis, N. Echols, J. J. Headd, L.-W. Hung, G. J. Kapral, R. W. Grosse-Kunstleve, A. J. McCoy, N. W. Moriarty, R. Oeffner, R. J. Read, D. C. Richardson, J. S. Richardson, T. C. Terwilliger and P. H. Zwart, Acta Crystallogr D Biol Crystallogr, 2010, 66, 213–221.

39 O. Trott and A. J. Olson, J Comput Chem, 2010, 31, 455–461.

40 K. Zinovjev and M. W. van der Kamp, Bioinformatics, 2020, 36, 5104–5106.

41 D. Case, R. Betz, D. S. Cerutti, T. Cheatham, T. Darden, R. Duke, T. J. Giese, H. Gohlke, Götz, N. Homeyer, S. Izadi, P. Janowski, J. Kaus, A. Kovalenko, T.-S. Lee, S. LeGrand, P. Li, C. Lin, T. Luchko and P. Kollman, Amber 16, University of California, San Francisco.

42 C. R. Søndergaard, M. H. M. Olsson, M. Rostkowski and J. H. Jensen, J Chem Theory Comput, 2011, 7, 2284–2295.

43 J. A. Maier, C. Martinez, K. Kasavajhala, L. Wickstrom, K. E. Hauser and C. Simmerling, J Chem Theory Comput, 2015, 11, 3696–3713.

44 J. Wang, R. M. Wolf, J. W. Caldwell, P. A. Kollman and D. A. Case, J Comput Chem, 2004, 25, 1157–1174.

45 S. Kumar, J. M. Rosenberg, D. Bouzida, R. H. Swendsen and P. A. Kollman, J Comput Chem, 1992, 13, 1011–1021.

46 B. R. Miller, T. D. McGee, J. M. Swails, N. Homeyer, H. Gohlke and A. E. Roitberg, J Chem Theory Comput, 2012, 8, 3314–3321.

47 D. R. Roe and T. E. Cheatham, J Chem Theory Comput, 2013, 9, 3084–3095.

