## Supplementary Information for "Delineation of the Complete Reaction Cycle of a Natural Diels-Alderase"

**This Supplement Includes:**

Table S1, pg 2

Figures S1 - S11, pgs 3 - 8

Equations S1 - S6, pg 9

Supplementary Methods and NMR spectra, pgs 10 - 22

Supplementary References, pg 23

### Supplementary Table

**Table S1.** Summary of X-ray data collection and refinement statistics.

| | AbyU-13 Complex |
|---|---|
| **Data collection** | |
| Beamline wavelength (Å) | 0.9999 |
| Space group | $P12_11$ |
| Cell dimensions | |
| *a, b, c* (Å) | 73.57, 61.56, 87.78 |
| a, b, g, (°) | 90.0, 112.96, 90.0 |
| Resolution (Å) | 61.56-1.95 (2.0-1.95)[a] |
| *Rpim* | 0.076 (0.938)[a] |
| No. of reflections | 332211 (24428)[a] |
| No. of unique reflections | 52793 (3710)[a] |
| $I/\sigma I$ | 7.2 (0.9)[a] |
| $CC_{1/2}$ (%) | 98.9 (29.8)[a] |
| Completeness (%) | 99.8 (100)[a] |
| Redundancy | 6.3 (6.6)[a] |
| **Refinement** | |
| $R_{\text{work}}/R_{\text{free}}$ | 0.218/0.245 |
| No. of atoms | |
| Protein | 4172 |
| Ligand/ion | 120 |
| Water | 42 |
| *B* factors Å$^2$ | |
| Protein | 50 |
| Ligand/ion | 76.7 |
| Water | 43.4 |
| Root mean square deviations | |
| Bond lengths (Å) | 0.007 |
| Bond angles (°) | 1.65 |
| Ramachandran favoured (%) | 95 |
| Ramachandran outliers (%) | 0.8 |

[a]Values in parentheses are for the highest resolution shell

### Supplementary Figures

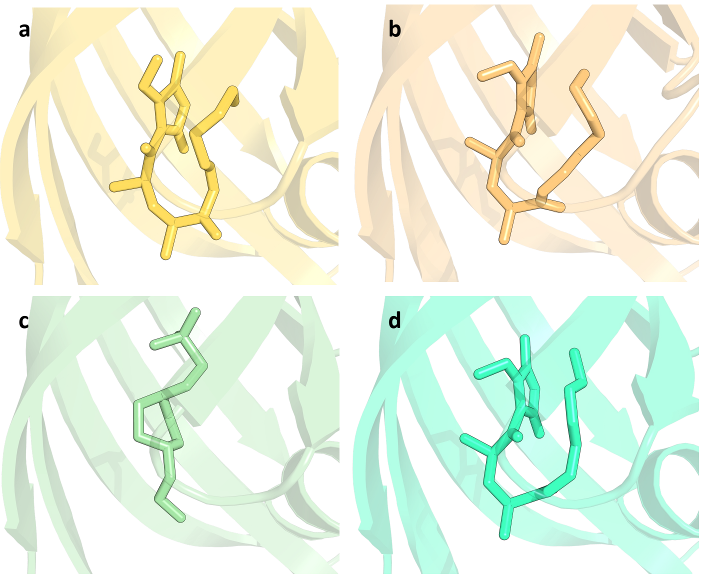


**Figure S1.** Active site occupancy in the four copies of AbyU (A-D) comprising the asymmetric unit. a) Chain A with **13**. b) Chain B with **13**. c) Chain C with HEPES. d) Chain D with **13**.

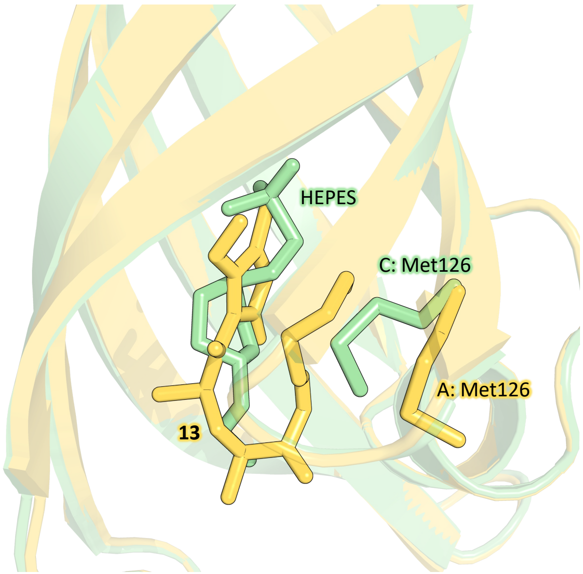


**Figure S2.** Comparison of the active sites of Chain A and Chain C. Superimposition of Chain A and Chain C from the asymmetric unit of 7PXO. Ligands and Met126 from each chain are shown in stick representation. Chain A is coloured pale green and Chain C is coloured pale orange.

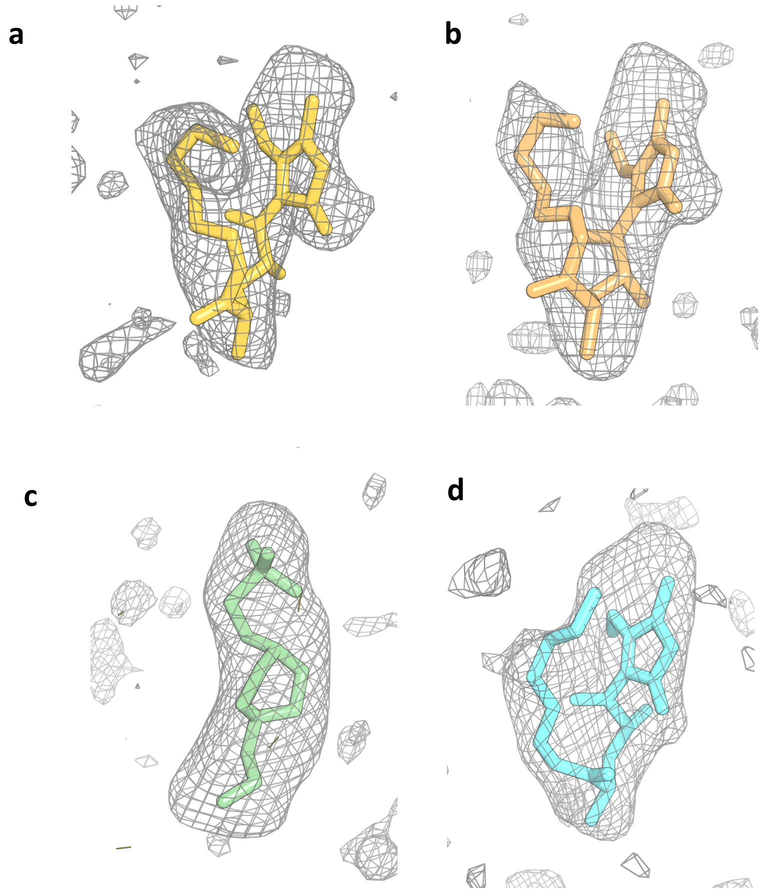


**Figure S3.** Polder OMIT maps of the molecules identified in the active sites of each the four molecules of AbyU that comprise the asymmetric unit, contoured at 3 $\sigma$. Ligands are shown in stick representation. a) **13** in Chain A, coloured yellow. b) **13** in Chain B, coloured pale orange. c) HEPES in Chain C, coloured pale green. d) **13** in Chain D, coloured turquoise.

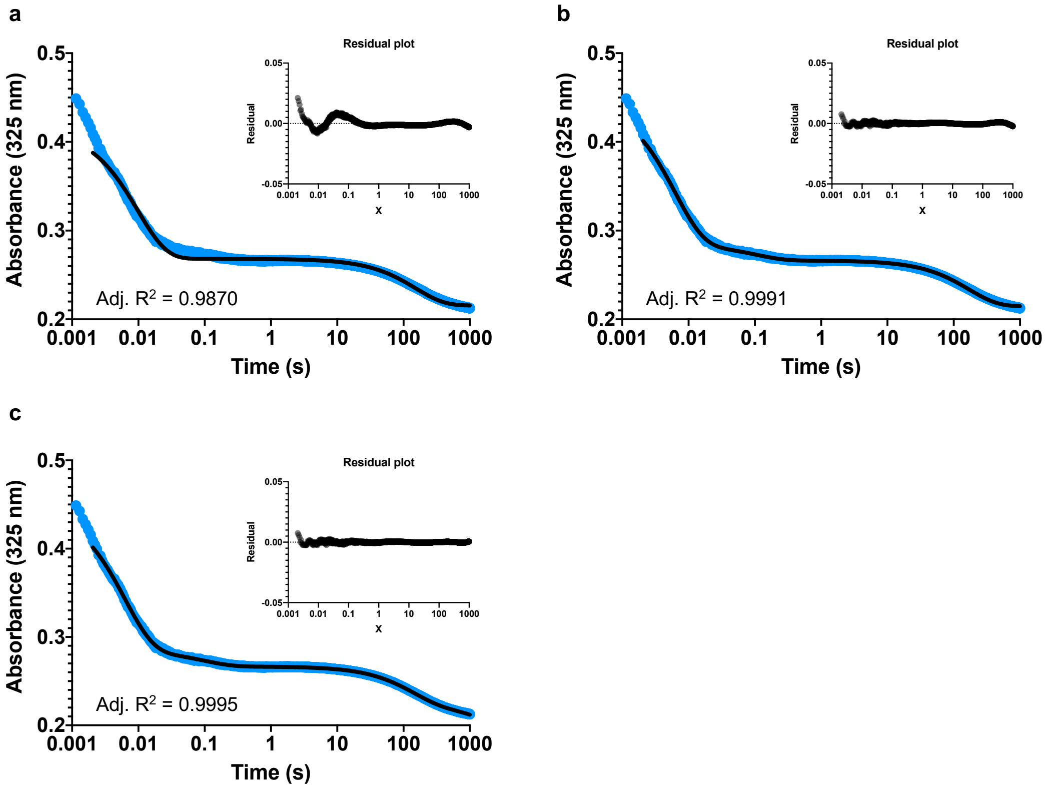


**Figure S4.** Fitting analysis for the reaction of 25 μM substrate **2** with 400 μM AbyU over 1000 seconds. The residual plots are inset and the adjusted $R^2$ value given below the transients. a) The reaction transient fitted to Equation S2, with two exponential terms. b) The reaction transient fitted to Equation 1, with three exponential terms. c) The reaction transient fitted to Equation S3 with four exponential terms.

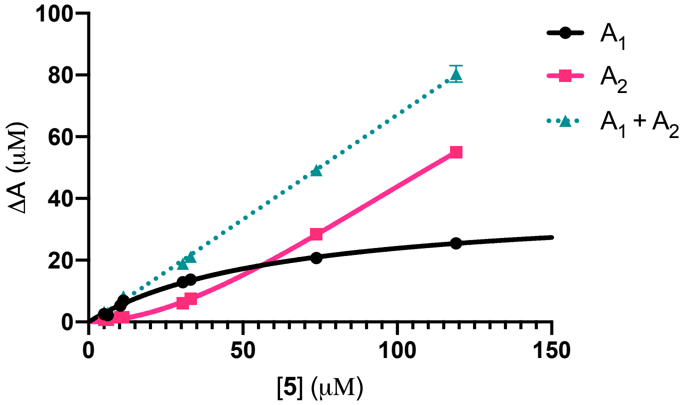


**Figure S5.** Analysis of the amplitude change observed during $k_1$ and $k_2$, denoted $A_1$ and $A_2$ respectively, with respect to substrate concentration. $A_1$ is fit to Equation S1 and $A_2$ is fit to equation S6, though this is qualitative due to an inability to saturate the curve. The quantity of substrate binding in the non-productive $k_3$ conformation can be determined by adding the amplitudes of $k_1$ and $k_2$, which fit to a straight line with the slope equal to 0.68 ± 0.006 and the y-intercept equal to -0.68.

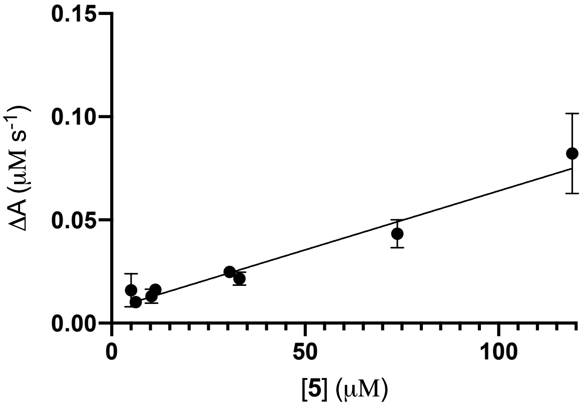


**Figure S6.** Concentration dependent analysis of $k_{obs3}$. Over 10 s this exponential curve is fit to a straight line and the rates plotted as a function of substrate concentration.

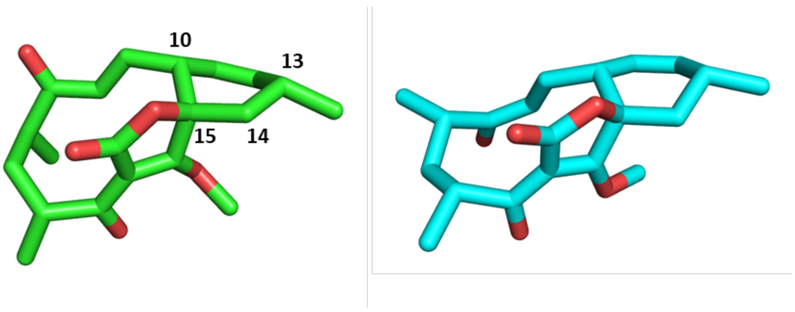


**Figure S7.** Possible reaction products of **5** - Abyssomicin C precursor analogue **6** (green carbon atoms) and its atropisomer **7** (cyan carbon atoms). Hydrogen atoms are not shown for clarity.

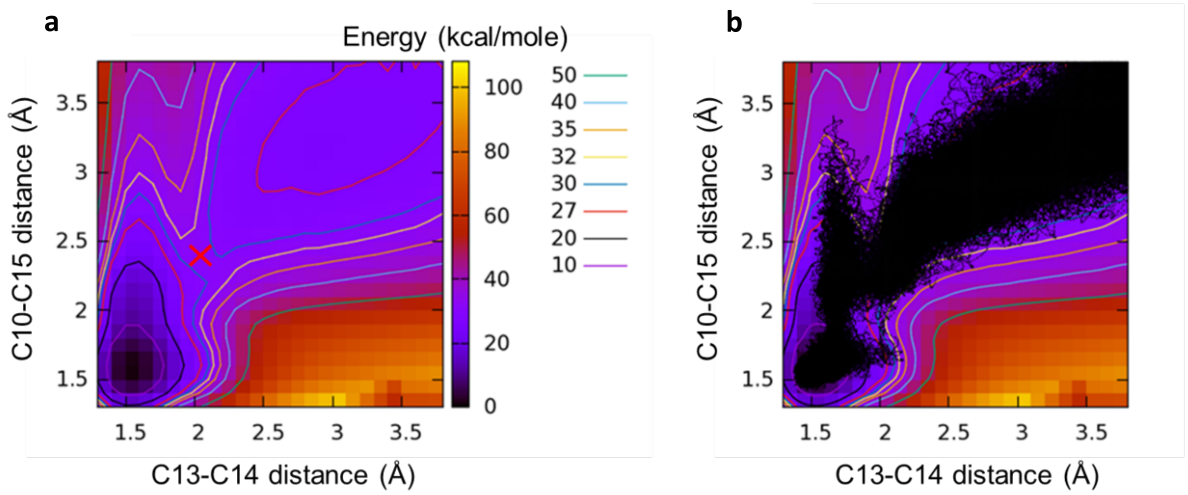


**Figure S8.** a) 2D potential energy surface at the SCC-DFTB/ff14SB level for the reaction of **7** in binding mode A in AbyU (see Figure 4 in the main manuscript and '2D Potential Energy Surface' section in Supplementary Methods, below). Approximate transition state location is indicated by a red cross. b) 2D potential energy surface with distances sampled during the 1D umbrella sampling reaction simulations superimposed (black dots). 1D sampling results shown for the 10 repeat umbrella sampling runs using the optimal reaction co-ordinate.

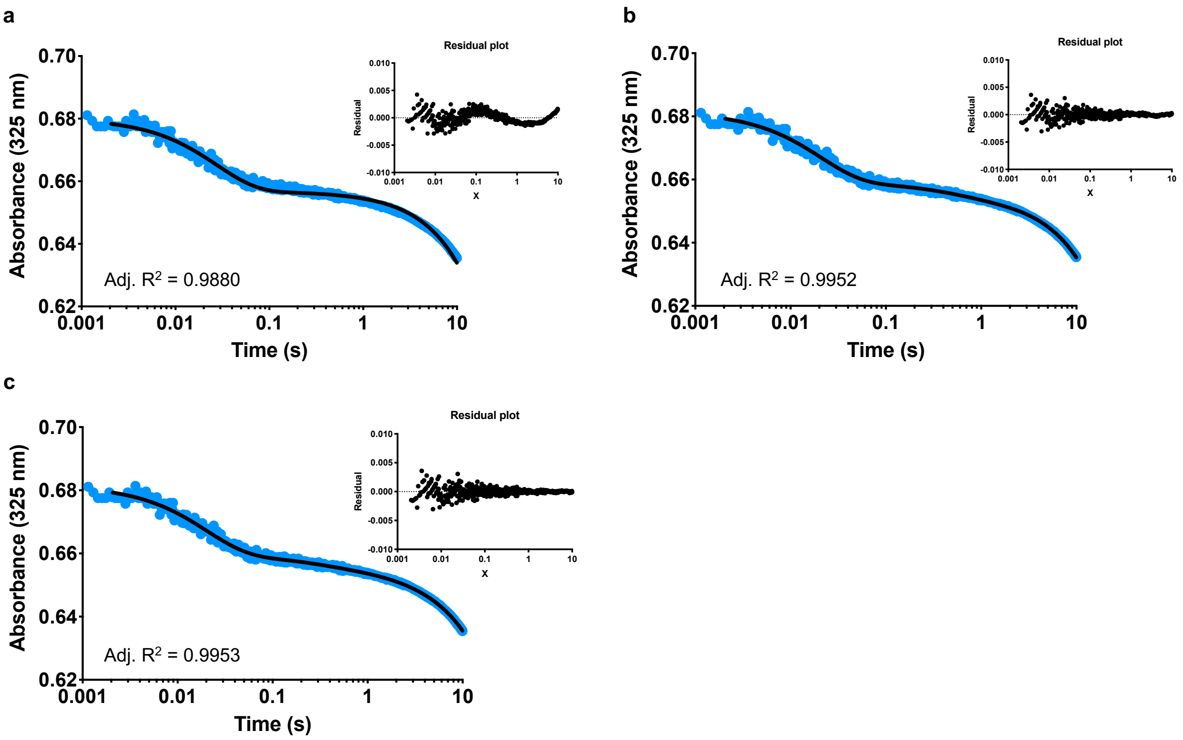


**Figure S9.** Fitting analysis for the reaction of 54 µM substrate **5** with 1 µM AbyU over 10 seconds. The residual plots are inset and the adjusted $R^2$ values are given below the transients. a) The reaction transient fitted to a single exponential and sloping baseline (Equation S4). b) The reaction transient fitted to Equation 3, with two exponential terms and a sloping baseline. c) The reaction transient fitted to Equation S5, with three exponential terms and a sloping baseline.

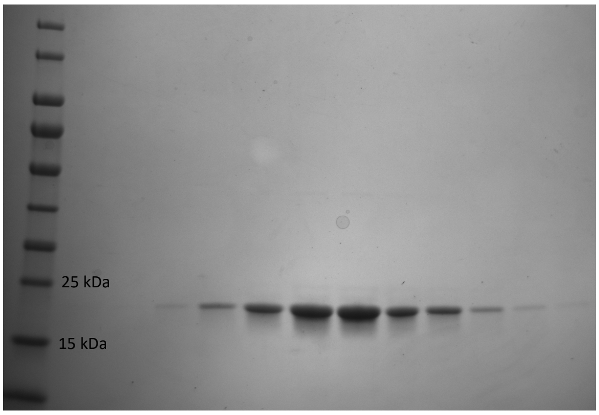


**Figure S10.** SDS-PAGE analysis of purified AbyU.

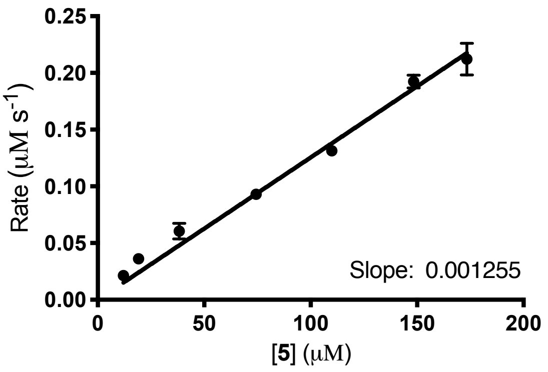


**Figure S11.** Rate of spontaneous cyclisation of **5** in buffer.

### Supplementary Equations

**Equation S1:**

$$\mathrm{Y} = \frac{V_{max} \times [\mathrm{S}]}{K_m + [\mathrm{S}]}$$

**Equation S2:**

$$\Delta A = \sum_{i=1}^{\leq 2} A_i \exp\left(-k_i t\right)$$

**Equation S3:**

$$\Delta A = \sum_{i=1}^{\leq 4} A_i \exp\left(-k_i t\right)$$

**Equation S4:**

$$\Delta A = A\exp(-kt) + mt$$

**Equation S5:**

$$\Delta A = \sum_{i=1}^{3} A_i \exp\left(-k_i t\right) + mt$$

**Equation S6:**

$$\mathrm{Y} = \frac{V_{max} \times [S]^h}{{k_{half}}^h + [S]^h}$$

**Methods**

1. **Chemical Synthesis**

**General Experimental**

All reactions were carried out using standard Schlenk syringe-septa techniques in flame dried glassware under a positive pressure of nitrogen in anhydrous solvents unless otherwise stated. Reagents and solvents were purchased from commercial suppliers and used without further purification unless reported. Anhydrous THF, $Et_2O$, hexane, DCM, toluene and MeCN were dried by passing through a modified Grubbs system of alumina columns and stored under nitrogen. MeOH, EtOH, EtOAc, DIPEA and TEA were dried by distillation from calcium hydride and stored under nitrogen and over 3 Å molecular sieves. Degassed solvents were prepared by freeze-pump-thaw cycling under nitrogen or by sparging with nitrogen.

Analytical thin layer chromatography (TLC) was carried out on Merck silica gel 60 $F_{254}$ analytical plates and were developed using UV fluorescence (254 nm) or $KMnO_4$ / Δ. Flash column chromatography was carried out on Sigma Aldrich silica gel 60 Å (43-63 μm) and an organic solvent system as stated. Infrared spectra were recorded on a Perkin-Elmer FT-IR spectrometer spectrum 2 with selected peaks of interested reported as absorption maxima ($cm^{-1}$). Mass spectrometry (MS) and High-resolution mass spectrometry (HRMS) were performed by the University of Bristol mass spectrometry service using electrospray ionisation (ESI) on a Bruker microOTOF II (TOF) or atmospheric pressure chemical ionisation (APCI) on a Thermo Scientific Orbitrap Elite (LC-Orbitrap). Optical rotation was measured on a Bellingham and Stanley Ltd. ADP220 polarimeter and is quoted in (° ml)(g dm)$^{-1}$. Melting points of solid products were recorded on a Stuart MP20.

NMR spectra were recorded on Varian 400-MR (400 MHz), Jeol ECS400 (400 MHz), Jeol ECZ400 (400 MHz), JeolVAR ECZ400 (400 MHz), BrukerNano400 (400 MHz), Bruker Avance III HD 500 Cryo (500 MHz), Bruker Neo 600 Cryo (600 MHz), and Bruker Avance III HD Cryo700 (700 MHz) spectrometers at ambient temperature. Spectra were recorded in deuterochloroform referenced to residual $CHCl_3$ ($^{1}H$, 7.26 ppm; $^{13}C$, 77.2 ppm), deuterated methanol referenced to residual MeOH ($^{1}H$, 3.30 ppm; $^{13}C$, 49.0 ppm) or deuterated acetone referenced to residual acetone ($^{1}H$, 2.05 ppm; $^{13}C$, 29.8 ppm). Chemical shifts (δ) are reported in parts per million (ppm) and coupling constants (*J*) are reported in Hertz (Hz). The following abbreviations are used to describe multiplicity: s (singlet), d (doublet), t (triplet), q (quartet), p (pentet), sext (sextet), br. (broad), ap. (apparent). COSY, HMBC, and HSQC NMR spectra were routinely used to definitively assign the signals of $^{1}H$ and $^{13}C$ NMR spectra. For clarity, the numbering of atoms does not correspond to the compound names.

### Chemical Synthesis of Substrate 5 and Product 6

Substrate **5** and Diels-Alder adduct **6** were synthesized according to the methods previously described.[1,2]

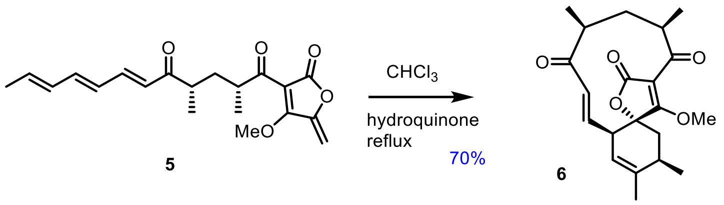


### (2*R*,4*S*,6*E*,8*E*,10*E*)-(5'-Methoxy-4'-methylene-2'-oxo-2',4'-dihydrofuran-1'-yl)-2,4-dimethyldodeca-6,8,10-triene-1,5-dione 5

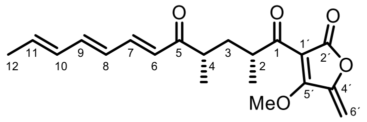


**5**

$\delta_H$ (400 MHz, $CDCl_3$) 7.25 (1H, overlapping m, 7-H), 6.58 (1H, dd, *J* 15.0, 10.5, 9-H), 6.30–6.11 (3H, m, 6-H, 8-H and 10-H), 5.96 (1H, m, 11-H), 5.26 (1H, d, *J* 3.0, 6'H*H*), 5.21 (1H, d, *J* 3.0, 6'-*H*H), 4.11 (3H, s, $OCH_3$), 3.64 (1H, m, 2-H), 2.81 (1H, m, 4-H), 2.21 (1H, m, 3-H*H*), 1.83 (3H, d, *J* 7.0, 12-$H_3$), 1.29 (1H, m, 3-*H*H), 1.15 (3H, d, *J* 6.5, 2-$CH_3$), 1.13 (3H, d, *J* 6.5, 4-$CH_3$). $\delta_C$ (100 MHz, $CDCl_3$) 203.4 (C-5), 200.7 (C-1), 168.7 (C-5'), 166.4 (C-2'), 148.9 (C-4'), 143.4 (C-7), 142.3 (C-9), 135.5 (C-11), 131.5 (C-10), 128.3 (C-8), 127.4 (C-6), 104.8 (C-1'), 95.9 (C-6'), 62.8 (O*C*$H_3$), 42.4 (C-4), 42.2 (C-4), 35.8 (C-3), 18.7 (C-12), 18.0 (2-$CH_3$), 17.1 (4-$CH_3$). All data are in accordance with the literature.[2]

### Diels Adduct 6

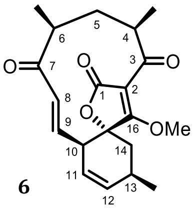


**6**

$\delta_H$ (500 MHz, $CDCl_3$) 6.47 (1H, dd, *J* 16.5, 6.0, 9-H), 6.25 (1H, d, *J* 16.5, 8-H), 5.86 (1H, *app*. dt, *J* 10.0, 3.0, 12-H), 5.68 (1H, *app.* dt, *J* 10.0, 3.0, 11-H), 3.91 (3H, s, OMe), 3.45 (1H, m, 10-H), 3.12 (1H, m, 4-H), 2.95 (1H, m, 6-H), 2.64 (1H, m, 13-H), 2.40 (1H, dd, *J* 14.5, 8.0, 14-*H*H), 1.87 (1H, ddd, *J* 15.5, 6.0, 4.0, 5-*H*H), 1.82 (1H, dd, *J* 14.5, 4.5, 14-H*H*), 1.21 (3H, d, *J* 7.0, 6-$CH_3$), 1.19 (3H, d, *J* 7.0, 4-$CH_3$), 1.17–1.13 (1H, overlapping m, 5-H*H*), 1.15 (3H, d, *J* 7.5, 13-$CH_3$). $\delta_C$ (125 MHz, $CDCl_3$) 204.3 (C-7), 200.6 (C-3), 178.2 (C-16), 169.9 (C-1), 141.5 (C-9), 136.7 (C-12), 131.6 (C-8), 121.8 (C-11), 107.0 (C-2), 86.1 (C-15), 61.7 ($OCH_3$), 46.65 (C-4), 46.60 (C-6), 44.6 (C-10), 39.0 (C-5), 36.6 (C-14), 29.3 (C-13) , 21.1 (13-$CH_3$), 17.0 (6-$CH_3$), 16.6 (4-$CH_3$). All data are in accordance with the literature.[2]

#### Synthesis of Non-transformable Substrate 13 for AbyU

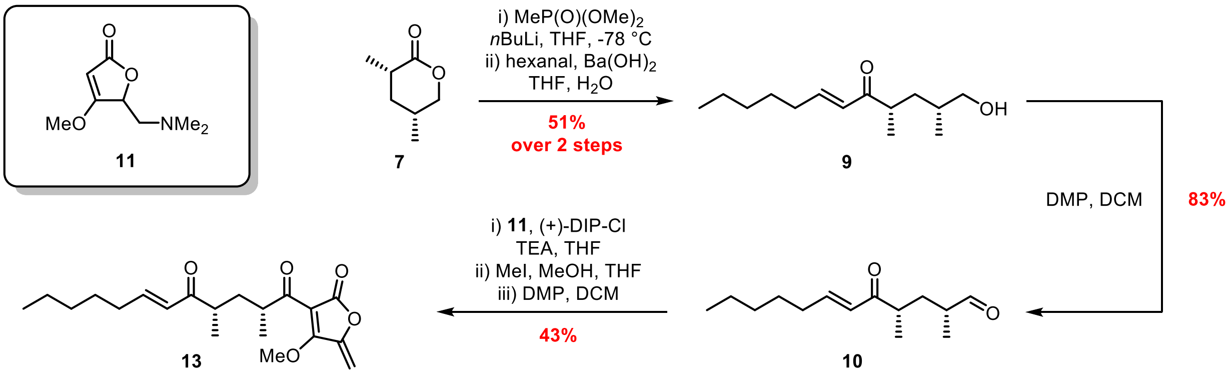


Lactone **7** and tetronate **11** were synthesised as previously described.[2]

**(2*R*,4*S*,*E*)-1-Hydroxy-2,4-dimethyldodec-6-en-5-one (9)**

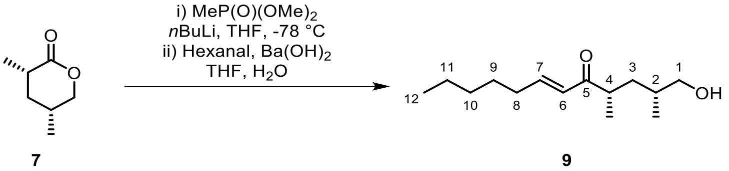


Dimethyl methylphosphonate (0.62 mL, 5.73 mmol) was dissolved in THF (10 mL) under nitrogen and cooled to -78 °C then *n*-BuLi in hexane (2.5 M, 2.29 mL, 5.73 mmol) was added dropwise over 5 minutes. The reaction mixture was stirred at -78 °C for 30 minutes then lactone **7** (350 mg, 2.73 mmol) in THF (5 mL) was added dropwise over 5 minutes. The reaction mixture was stirred at -78 °C for 1 hour the quenched with aqueous saturated $NH_4Cl$ (20 mL) and warmed to room temperature. The resulting solution was diluted with water (20 mL) and extracted with EtOAc (5 × 75 mL). The combined organic layers were dried over $Na_2SO_4$ and the solvent removed *in vacuo*. The crude material was dissolved in THF (55 mL) and cooled to 0 °C then $Ba(OH_2)$ (702 mg, 4.10 mmol) and hexanal (0.50 mL, 4.10 mmol) were added sequentially. Water (9 mL) was added dropwise and the reaction mixture was stirred at room temperature for 18 hours then quenched with aqueous saturated $NH_4Cl$ (50 mL). The resulting solution was extracted with EtOAc (3 × 100 mL), then the combined organic layers were dried over $Na_2SO_4$ and the solvent removed *in vacuo*. The crude material was purified by flash column chromatography (20-30% EtOAc in petroleum ether 40-60 °C + 0.05% TEA) to afford alcohol **9** (0.32 g, 51% over two steps) as a colourless oil; $[\alpha]_D^{25}$ = +8.0 (*c* 0.5, $CHCl_3$); $\nu_{max}$ (film) 3444, 2958, 2929, 2873, 2859, 1691, 1668, 1625, 1460; $\delta_H$ (400 MHz, $CO(CD_3)_2$) 6.90 (1H, dt, *J* 15.7, 7.0, 7-H), 6.20 (1H, dt, *J* 15.7, 1.5, 6-H), 3.42 – 3.36 (1H, m, 1-*H*H), 3.34 – 3.27 (1H, m, 1-H*H*), 2.98 (1H, ap. dp, *J* 8.3, 6.9, 4-H), 2.23 (2H, ap. qd, *J* 7.0, 1.5, 8-$H_2$), 1.81 (1H, ddd, *J* 14.0, 8.3, 6.1, 3-*H*H), 1.57 – 1.43 (3H, m, 2-H and 9-$H_2$), 1.36 – 1.27 (4H, m, 10-$H_2$ and 11-$H_2$), 1.09 – 1.05 (1H, m, 3-H*H*), 1.04 (3H, d, *J* 6.9, 4-$CH_3$), 0.92 – 0.86 (6H, m, 2-$CH_3$ and 12-$H_3$); $\delta_C$ (101 MHz, $CO(CD_3)_2$) 203.5 (C-5), 147.5 (C-7), 130.0 (C-6), 67.7 (C-1), 41.7 (C-4),

37.9 (C-3), 34.5 (C-2), 32.9 (C-8), 32.1 (C-10), 28.6 (C-9), 23.0 (C-11), 17.9 ($CH_3$-4), 17.5 (2-$CH_3$), 14.2 (C-12); HRMS (ESI) calc. for $[C_{14}H_{26}O_2]^+$ 227.2006 Found 227.1999.
$\delta_H$ (400 MHz, $CD_3OD$) 6.97 (1H, dt, *J* 15.8, 7.0, 7-H), 6.22 (1H, dt, *J* 15.8, 1.5 Hz, 6-H), 3.39 (1H, dd, *J* 10.7, 5.5, 1-*H*H), 3.30 (1H, dd, *J* 10.7, 6.4, 1-H*H*), 3.02 (1H, dqd, *J* 8.7, 6.9, 5.6, 4-H), 2.26 (2H, ap. qd, *J* 7.0, 1.5, 8-$H_2$), 1.81 (1H, ddd, *J* 13.9, 8.7, 5.6, 3-*H*H), 1.56 – 1.46 (3H, m, 2-H and 9-$H_2$), 1.38 – 1.30 (4H, m, 10-$H_2$ and 11-$H_2$), 1.14 – 1.10 (1H, m, 3-H*H*), 1.08 (3H, d, *J* 6.9, 4-$CH_3$), 0.94 – 0.88 (6H, m, 2-$CH_3$ and 12-$H_3$).
NB alcohol **9** undergoes cyclisation to the lactol in both $CDCl_3$ and $CD_3OD$.

#### (2*R*,4*S*,*E*)-2,4-Dimethyl-5-oxododec-6-enal (10)

Method 1 – Oxidation with TEMPO/BAIB:

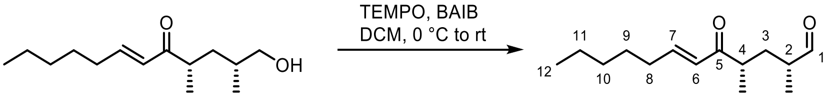


**9** **10**

Alcohol **9** (240 mg, 1.06 mmol) was dissolved in DCM (5.5 mL) under nitrogen then cooled to 0 °C. BAIB (820 mg, 2.54 mmol) and TEMPO (66 mg, 0.42 mmol) were added sequentially, and the reaction mixture was stirred at room temperature for 45 minutes. The reaction mixture was quenched with aqueous saturated $Na_2S_2O_3$ (4 mL) and aqueous saturated $NaHCO_3$ (1 mL) then stirred vigorously for 1 hour. The resulting solution was diluted with DCM (30 mL) and water (20 mL) and the layers separated. The aqueous layer was extracted with further DCM (2 x 50 mL) and the combined organic layers were washed with brine (50 mL), dried over $MgSO_4$ and the solvent removed *in vacuo*. The crude material was purified by flash column chromatography (90% DCM and 1% $Et_2O$ in petroleum ether 40-60 °C) to afford aldehyde **10** (90 mg, 38%) as a colourless oil; $[\alpha]_D^{22}$ = +16.0 (*c* 0.1, $CHCl_3$); $\nu_{max}$ (film) 2962, 2931, 2874, 2859, 1726, 1694, 1668, 1626, 1214, 750; $\delta_H$ (400 MHz, $CDCl_3$) 9.59 (1H, d, *J* 1.8, 1-H), 6.91 (1H, dt, *J* 15.7, 6.9, 7-H), 6.15 (1H, dt, *J* 15.7, 1.5, 6-H), 2.91 – 2.81 (1H, m, 4-H), 2.37 (1H, sextetd, *J* 7.0, 1.8, 2-H), 2.26 – 2.12 (3H, m, 3-*H*H and 8-$H_2$), 1.52 – 1.40 (2H, m, 9-$H_2$), 1.36 – 1.24 (5H, m, 3-H*H*, 10-$H_2$ and 11-$H_2$), 1.12 (3H, d, *J* 7.0, 4-$CH_3$), 1.08 (3H, d, *J* 7.0, 2-$CH_3$), 0.93 – 0.86 (3H, m, 12-$H_3$); $\delta_C$ (101 MHz, $CDCl_3$) 204.5 (C-1), 203.1 (C-5), 148.5 (C-7), 128.8 (C-6), 44.4 (C-2), 41.1 (C-4), 33.5 (C-3), 32.6 (C-8), 31.5 (C-10), 27.9 (C-9), 22.5 (C-11), 17.6 ($CH_3$-4), 14.1 (C-12), 13.9 ($CH_3$-2); HRMS (ESI) calc. for $[C_{14}H_{24}O_2]^+$ 225.1849 Found 225.1843.

Method 2 – Oxidation with DMP:

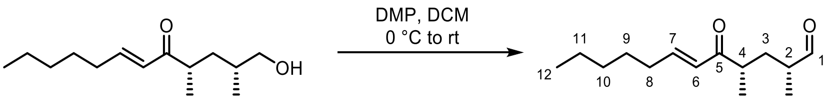


**9** **10**

Alcohol **9** (200 mg, 0.88 mmol) was dissolved in DCM (9 mL) under nitrogen and cooled to 0 °C, then DMP (600 mg, 1.41 mmol) was added in one portion. The reaction mixture was stirred at room temperature for 30 minutes then quenched with aqueous saturated $Na_2S_2O_3$ (3 mL) and aqueous saturated $NaHCO_3$ (1 mL). The resulting solution was diluted with water (20 mL) and extracted with DCM (3 x 30 mL). The combined organic layers were dried over $MgSO_4$ and the solvent removed *in vacuo*. The crude material was purified by flash column chromatography (20% $Et_2O$ in petroleum ether 40-60 °C) to afford **10** (164 mg, 83%) as a colourless oil. Data consistent with that reported above.

**(2*R*,4*S*,*E*)-1-(4'-Methoxy-5'-methylene-2'-oxo-2',5'-dihydrofuran-3'-yl)-2,4-dimethyldodec-6-ene-1,5-dione (13)**

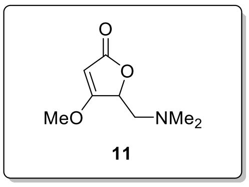


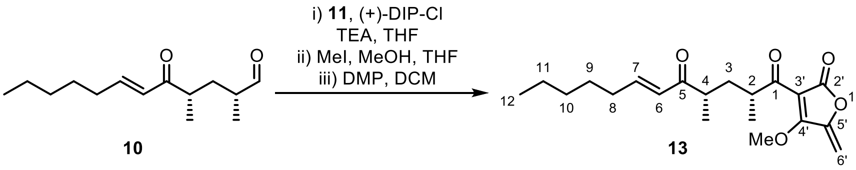


Tetronate **11** (143 mg, 0.83 mmol) was dissolved in THF (2.7 mL) under nitrogen and cooled to -78 °C, then TEA (0.22 mL, 1.60 mmol) followed by (+)-DIP-Cl (0.74 M in hexane, 1.13 mL, 0.83 mmol) were added dropwise. The reaction mixture was stirred for 30 minutes at -78 °C then aldehyde **10** (85 mg, 0.38 mmol) in THF (1.3 mL) was added dropwise and the resulting solution was stirred for a further hour at -78 °C. The reaction mixture was warmed to 0 °C then MeOH (0.42 mL) and MeI (0.24 mL, 3.80 mmol) were added sequentially. The reaction mixture was stirred for 3 hours at 0 °C then quenched with aqueous saturated $NaHCO_3$ (2.5 mL) and the resulting solution stirred vigorously at room temperature for 1 hour. The reaction mixture was diluted with water (20 mL) and extracted with EtOAc (3 x 30 mL). The combined organic layers were dried over $MgSO_4$ and the solvent concentrated *in vacuo* to ~2 mL, then DCM (20 mL) was added and the resulting solution concentrated *in vacuo* to ~1 mL. The crude material was purified by flash column chromatography (40% $Et_2O$ in petroleum ether 40-60 °C) to afford a mixture of epimeric alcohols which were used immediately in the next step. NB polymerisation of the intermediate alcohol occurs rapidly upon concentration therefore the intermediate was never fully dried.

The mixture of alcohol epimers was dissolved in DCM (4 mL) under nitrogen and cooled to 0 °C, then DMP (161 mg, 0.38 mmol) was added in one portion. The reaction mixture was stirred at room temperature for 45 minutes then quenched with aqueous saturated $Na_2S_2O_3$ (3 mL) and aqueous saturated $NaHCO_3$ (1 mL). The resulting solution was diluted with water (20 mL) and extracted with DCM (3 x 30 mL). The combined organic layers were dried over $MgSO_4$ and the solvent removed *in vacuo*. The crude material was purified by flash column chromatography (10% EtOAc in petroleum ether 40-60 °C) to afford **13** (57 mg, 43%) as a yellow oil; $[\alpha]_D^{22}$ = -16.0 (*c* 0.1, $CHCl_3$); $\nu_{max}$ (film) 2959, 2931, 2873, 2858, 1770, 1687, 1666, 1590, 748; $\delta_H$ (400 MHz, $CDCl_3$) 6.90 (1H, dt, *J* 15.8, 6.9, 7-H), 6.14 (1H, dt, *J* 15.8, 1.5, 6-H), 5.26 (1H, d, *J* 2.8, 6'-*H*H), 5.21 (1H, d, *J* 2.8, 6'-H*H*), 4.11 (3H, s, $OCH_3$), 3.62 (1H, dqd, *J* 8.2, 7.0, 5.6, 2-H), 2.82 (1H, ap. sext, *J* 7.0, 4-H), 2.26 – 2.13 (3H, m, 3-*H*H and 8-$H_2$), 1.52 – 1.39 (2H, m, 9-$H_2$), 1.36 – 1.20 (5-H, m, 3-H*H*, 10-$H_2$ and 11-$H_2$), 1.14 (3H, d, *J* 7.0, 2-$CH_3$), 1.11 (3H, d, *J* 7.0, 4-$CH_3$), 0.90 – 0.85 (3H, m, 12-$H_3$); $\delta_C$ (101 MHz, $CDCl_3$) 203.5 (C-5), 200.7 (C-1), 168.7 (C-5'), 166.4 (C-2'), 149.0 (C-4'), 148.2 (C-7), 128.9 (C-6), 104.8 (C-3'), 96.0 (C-6'), 62.8 ($OCH_3$), 42.4 (C-2), 41.6 (C-4), 35.7 (C-3), 32.7 (C-8), 31.5 (C-10), 27.9 (C-9), 22.6 (C-11), 18.0 ($CH_3$-2), 17.1 ($CH_3$-4), 14.1 (C-12); HRMS (ESI) calc. for $[C_{20}H_{28}O_5]^+$ 349.2010 Found 349.2001.

### NMR Spectra

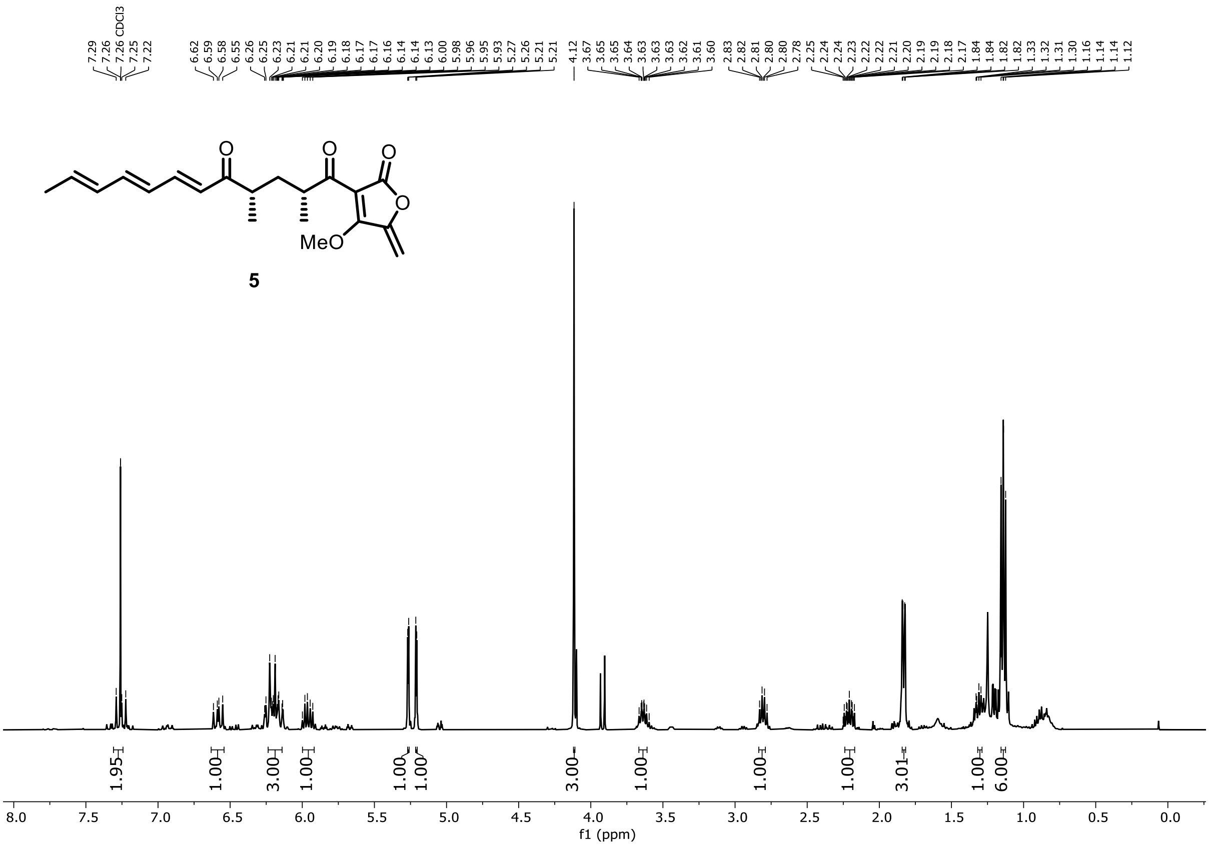


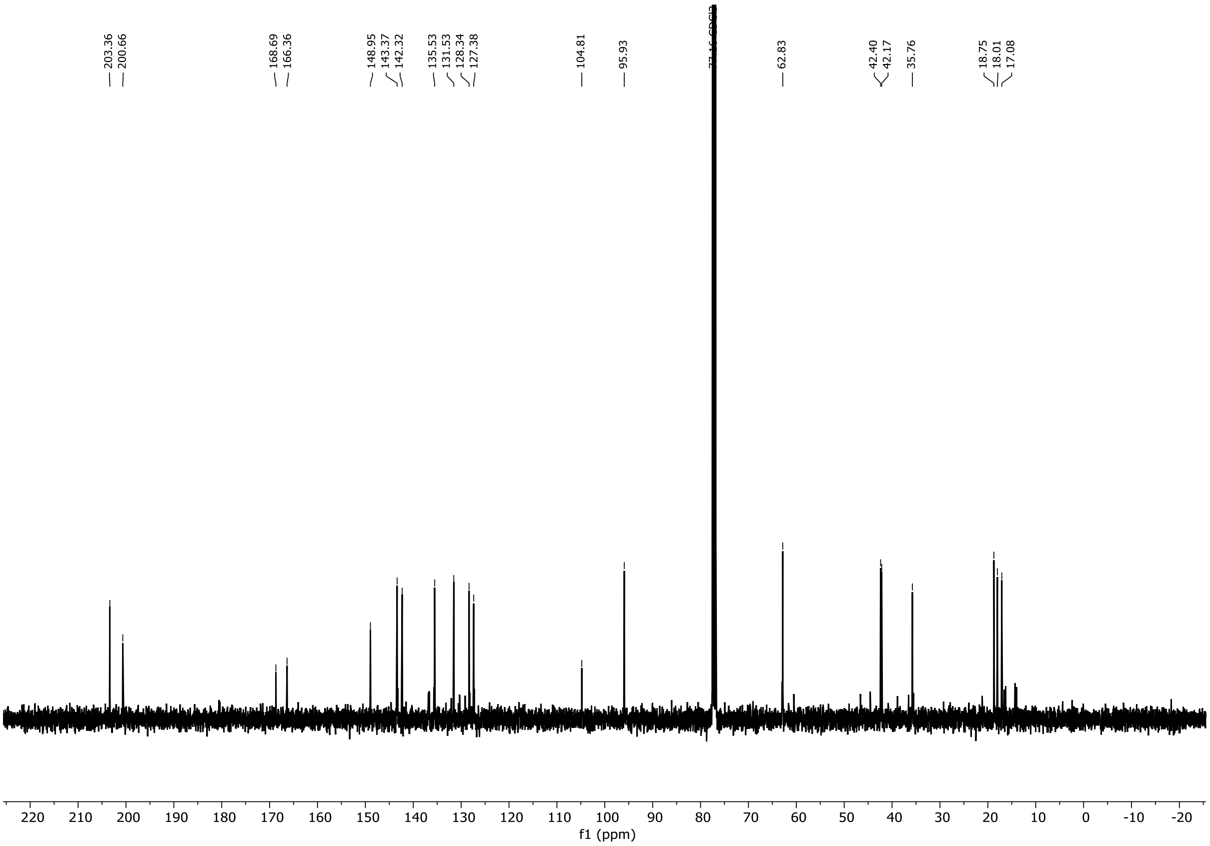

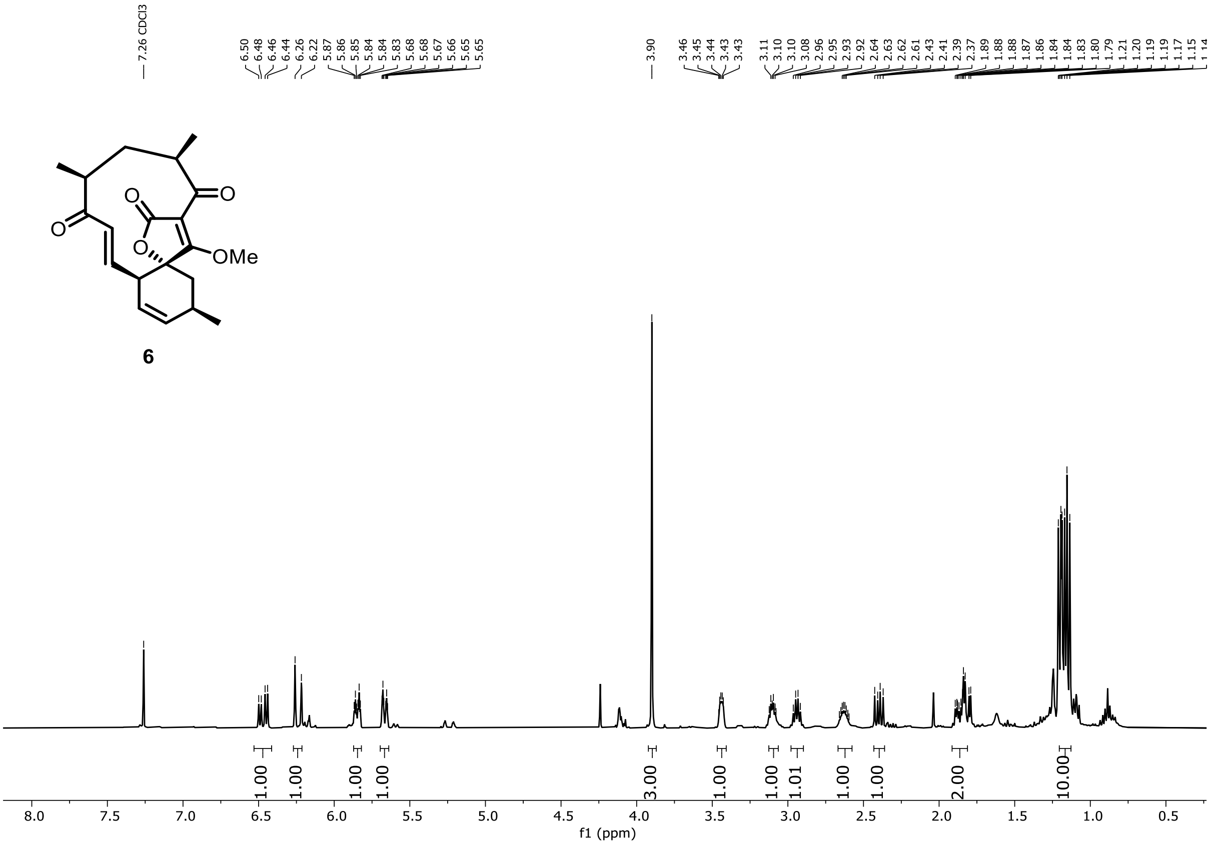
7.26 CDCl3
6.50 6.48 6.46 6.44 6.26 6.22 5.87 5.86 5.85 5.84 5.84 5.83 5.68 5.68 5.67 5.66 5.65 5.65
3.90
3.46 3.45 3.44 3.43 3.43
3.11 3.10 3.10 3.08 2.96 2.95 2.93 2.92 2.64 2.63 2.62 2.61 2.43 2.41 2.39 2.37 1.89 1.88 1.88 1.87 1.86 1.84 1.84 1.83 1.80 1.79 1.21 1.20 1.19 1.19 1.17 1.15
O
O
O
O
OMe
6
1.00 1.00 1.00 1.00 3.00 1.00 1.00 1.01 1.00 1.00 2.00 10.00
8.0 7.5 7.0 6.5 6.0 5.5 5.0 4.5 4.0 3.5 3.0 2.5 2.0 1.5 1.0 0.5
f1 (ppm)

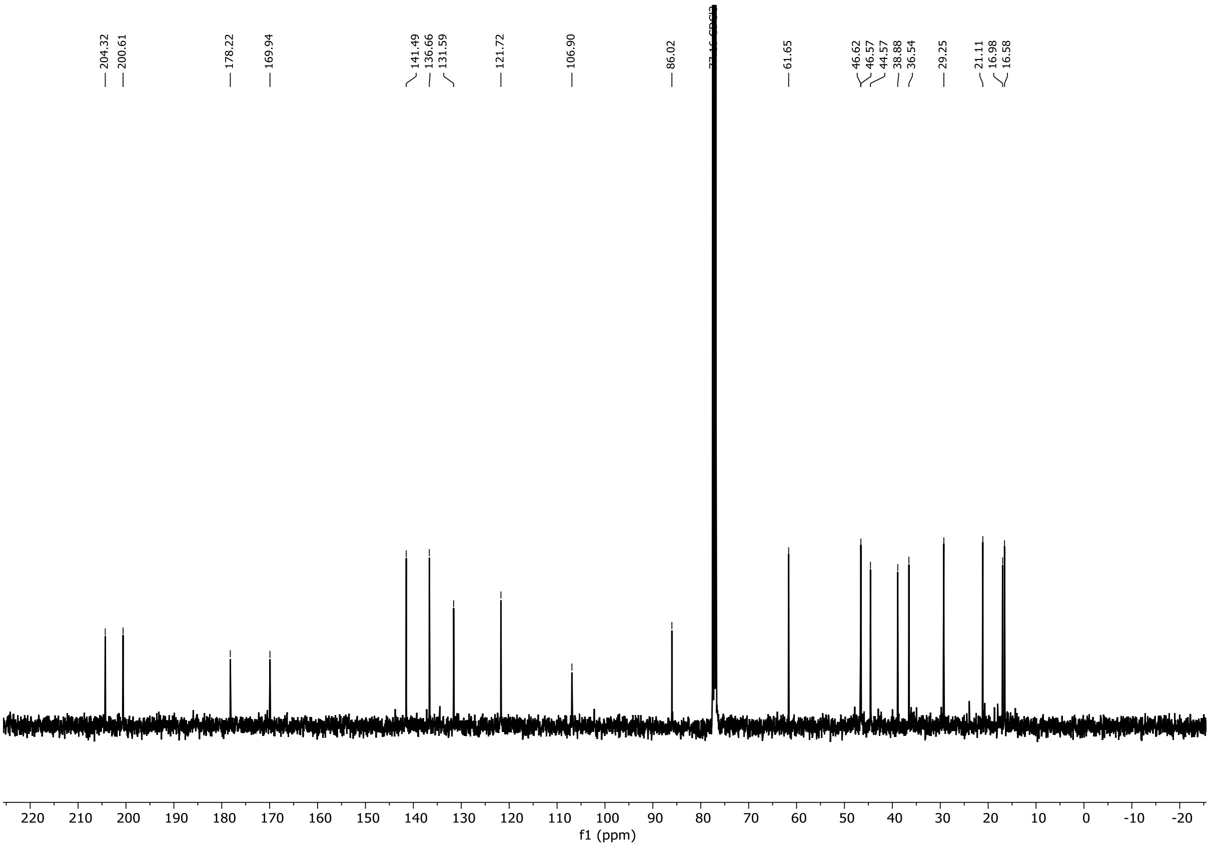
204.32 200.61 178.22 169.94 141.49 136.66 131.59 121.72 106.90 86.02 61.65 46.62 46.57 44.57 38.88 36.54 29.25 21.11 16.98 16.58
220 210 200 190 180 170 160 150 140 130 120 110 100 90 80 70 60 50 40 30 20 10 0 -10 -20
f1 (ppm)

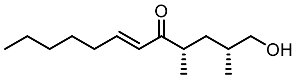

**9**

$^{1}H$-NMR in $(CD_3)_2CO$

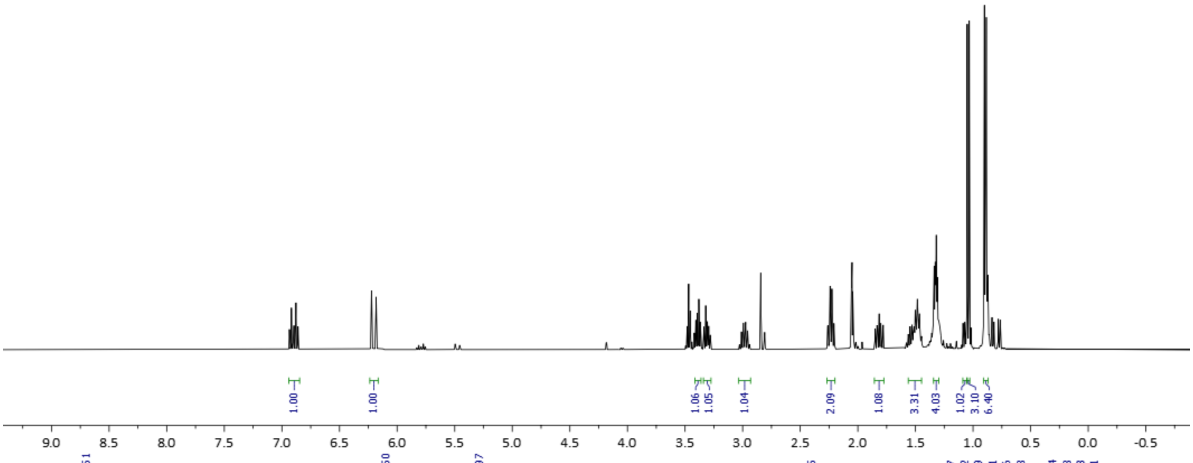


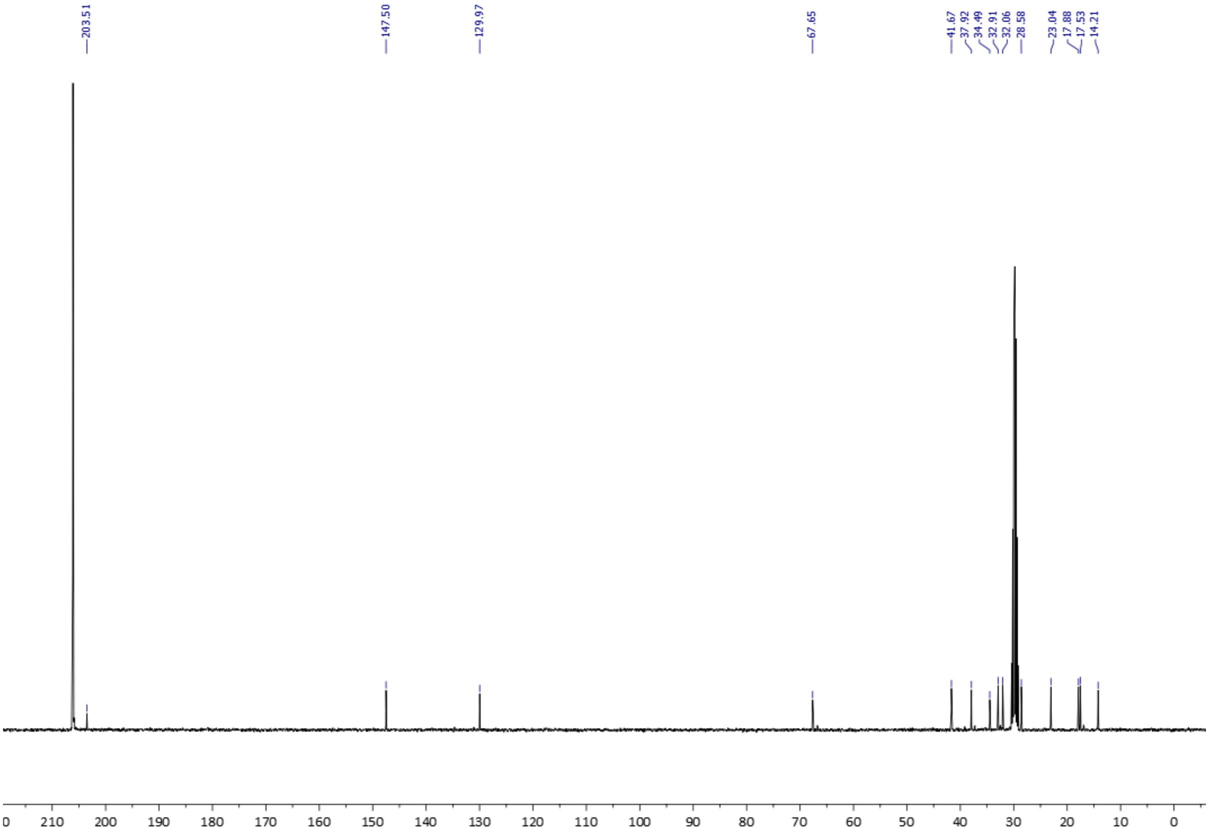

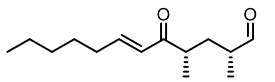

**10**

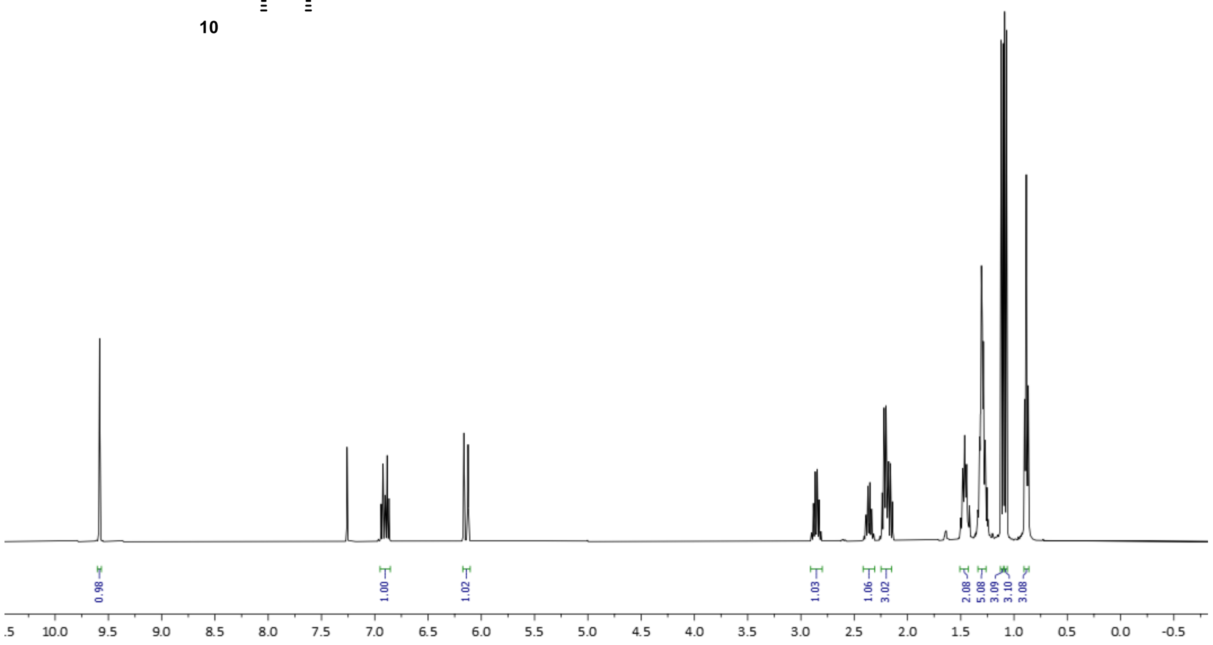
0.98
1.00
1.02
1.03
1.06
3.02
2.08
5.08
3.09
3.10
3.08
.5 10.0 9.5 9.0 8.5 8.0 7.5 7.0 6.5 6.0 5.5 5.0 4.5 4.0 3.5 3.0 2.5 2.0 1.5 1.0 0.5 0.0 -0.5

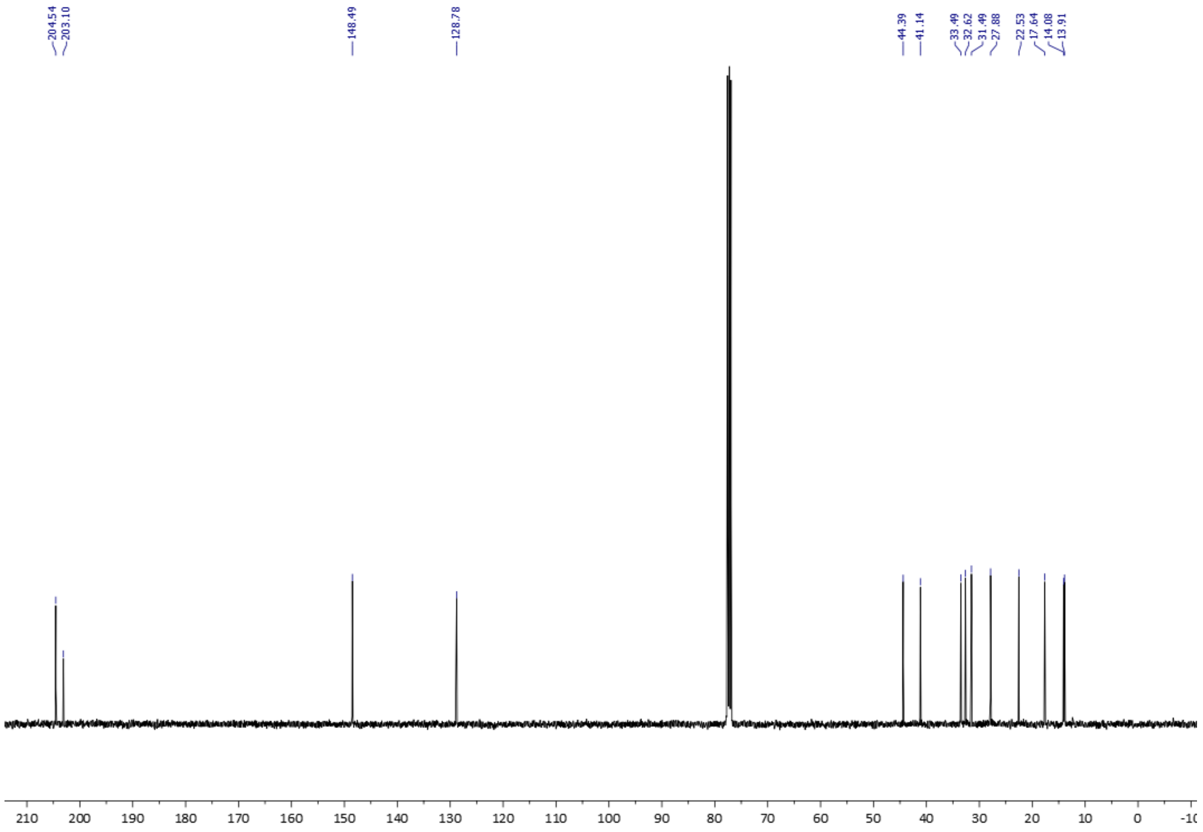
204.54
203.10
148.49
128.78
44.39
41.14
33.49
32.62
31.49
27.88
22.53
17.64
14.08
13.91
210 200 190 180 170 160 150 140 130 120 110 100 90 80 70 60 50 40 30 20 10 0 -1(

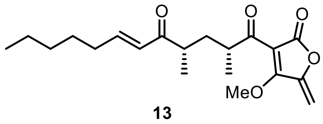
MeO

**13**

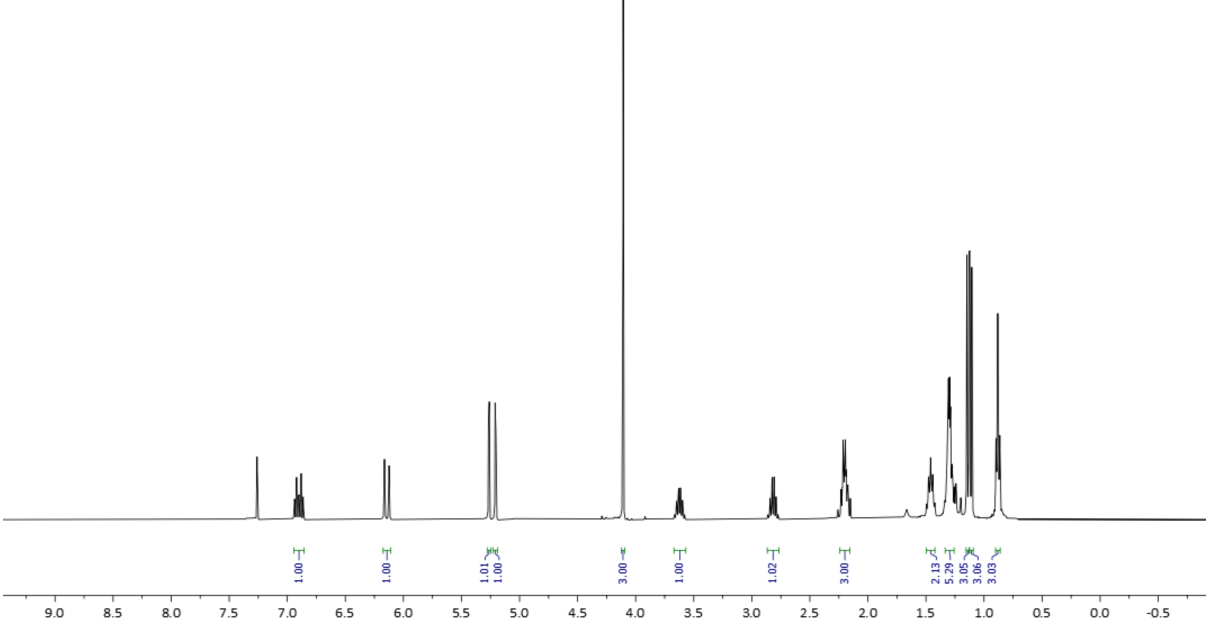
1.00
1.00
1.01
1.00
3.00
1.00
1.02
3.00
2.13
5.29
3.05
3.06
3.03
9.0 8.5 8.0 7.5 7.0 6.5 6.0 5.5 5.0 4.5 4.0 3.5 3.0 2.5 2.0 1.5 1.0 0.5 0.0 -0.5

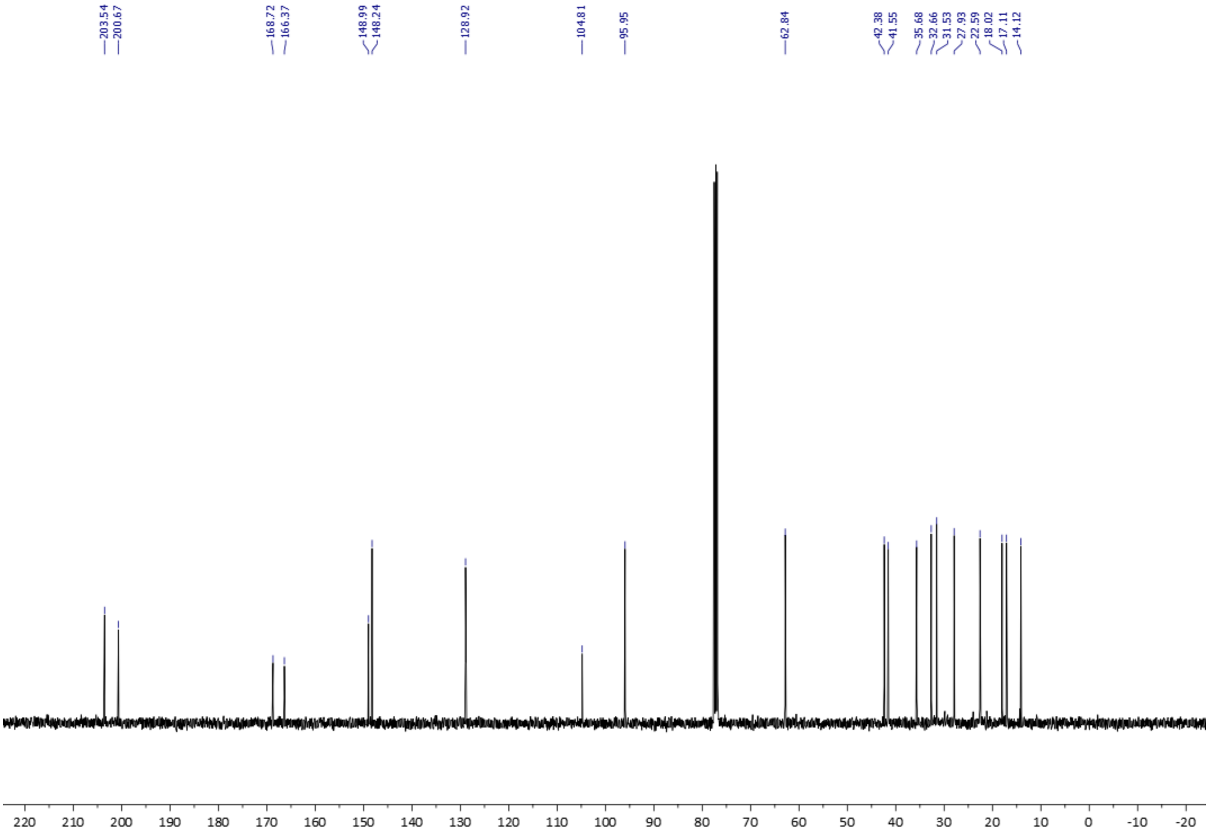
203.54
200.67
168.72
166.37
148.99
148.24
128.92
104.81
95.95
62.84
42.38
41.55
35.68
32.66
31.53
27.93
22.59
18.02
17.11
14.12
220 210 200 190 180 170 160 150 140 130 120 110 100 90 80 70 60 50 40 30 20 10 0 -10 -20

### 2. Gene cloning, expression and purification of AbyU

AbyU was cloned, expressed and purified according to the protocol outlined by Byrne *et al.*[1] Briefly, AbyU was cloned into the plasmid pOPINF and subsequently transformed into *E. coli* BL21 (DE3) cells for protein expression. Transformants were used to inoculate starter cultures supplemented with 100 µg/mL carbenicillin, which were in turn grown to turbidity overnight. These were used as a 1 % inoculant for 1 L of LB broth in a baffled flask, also supplemented with 100 µg/mL carbenicillin. Once an $OD_{600\ nm}$ of ~0.6 had been reached cultures were induced with the addition of IPTG to a final concentration of 1 mM. Cultures were grown overnight at 20 °C with shaking and cells harvested by centrifugation. Cell pellets were lysed using a cell disruptor (Constant Systems, UK) and clarified by centrifugation. AbyU was subsequently purified from the lysate by nickel affinity chromatography and size exclusion chromatography. The final elution step was performed in SEC buffer (20 mM Tris-HCl, 150 mM NaCl, pH 7.5). Fractions determined to be of > 95 % purity were pooled and concentrated using a Vivaspin 20 mL centrifugal concentrator (Sartorius).

### 3. Potential Energy Surface

To obtain the potential energy surface for the enzymatic reaction at the SCC-DFTB/ff14SB level, a series of energy minimizations is performed, ensuring no jumps to different potential energy surfaces occur. This was performed for product **7** starting from binding mode A from docking. To prepare the system, the relevant docked complex structure output after the reduce stage of the PREP protocol was taken and the remaining preparation steps were performed as before, with the exception that the system was instead solvated with a periodic box of TIP3P (minimal distance between the protein and the edge of the box 11 Å, 'closeness' parameter 0.75) and neutralized with 4 $Na^+$ counterions (this is to circumvent issues with the minimization algorithms in Amber16 when using a solvation sphere only). Then, to obtain a reasonable starting structure for obtaining the 2D potential energy surface, the system was briefly equilibrated with the following procedure, treating the whole structure MM (all steps performed with the Amber16 program pmemd.MPI, using periodic boundary conditions, a cut-off for direct-space non-bonded interactions of 8 Å and Particle-mesh Ewald summation for electrostatic interactions outside the cut-off):

- optimize solvent and solute hydrogen positions (minimization for 300 steps with restraints on solute heavy atoms, force constant: 100 kcal $mol^{-1}Å^{-2}$);
- heating/equilibration of solvent (initial random velocities assigned at 50K followed by 50 ps NPT heating to 300K with a 2 fs timestep using SHAKE restraints, using Langevin

dynamics with collision frequency 2 and the Berendsen barostat with a pressure relaxation time of 1 ps. All solute atoms were restrained with force constant 25 kcal $mol^{-1}Å^{-2}$);

- 300 steps of minimization and quick heating to 298 K (20 ps in NVT ensemble with 2 fs timestep, using Langevin dynamics with collision frequency 1, SHAKE and initial random velocities assigned at 25K);
- 300 ps equilibration in the NPT ensemble (Langevin dynamics with 2 fs timestep, SHAKE and collision frequency of 1 and using the Berendsen barostat with a pressure relaxation time of 1 ps).

Following this, the product and the sidechain of Trp124 were treated with SCC-DFTB. The structure at the end of the 300 ps equilibration was first minimized for 500 steps using steepest descent/conjugate gradient and then further minimized with the LBFGS method with a convergence criterion of 0.002 $mol^{-1}$ $Å^{-1}$ for energy gradients (all calculations performed using sander in AmberTools16).

To get the full energy surface, an initial scan was first performed along a diagonal path, with both bond distances restrained to the same value. To do this, optimizations were performed (again using the LBFGS method with a convergence criterion of 0.002 kcal $mol^{-1}Å^{-1}$) harmonically restraining the C10-C15 and C13-C14 bond distances (force constant 2500 kcal $mol^{-1}Å^{-2}$) to increasingly larger distances. The first structure was optimized with distances restrained to 1.3 Å and then the following structures were optimized in 0.1 Å steps, using the previous optimized structure as the starting structure, up to a distance of 3.8 Å. This path was scanned back and forth until energies converged to a smooth profile. During the optimizations, only residues that had at least one atom within 5 Å of the ligand were allowed to move freely. All other residues were restrained to the starting coordinates by a 50 kcal $mol^{-1}Å^{-2}$ harmonic potential. This was to ensure no discontinuities occur between progressive energies due to structural changes unrelated to the reaction. Separate scans were then performed along the C10-C15 distance starting from each optimized structure from the previous diagonal scan, keeping the C13-C14 distance restrained to the same value. Again, structures were optimized in 0.1 Å steps, starting from the initial structure and either increasing the C10-C15 distance to 3.8 Å, or decreasing it to 1.3 Å, to cover the entire surface. In this case only a single scan was performed in either direction as this already resulted in a smooth energy surface (except for part of one edge, which is unimportant due to its high relative energies).

**Supplementary References**

(1) Byrne, M. J.; Lees, N. R.; Han, L.-C.; Kamp, M. W. van der; Mulholland, A. J.; Stach, J. E.; Willis,C. L.; Race, P. R. *J. Am. Chem. Soc.* 2016, **138**, 6095–6098.

(2) A. J. Devine, A. E. Parnell, C. R. Back, N. R. Lees, S. T. Johns, A. Z. Zulkepli, R. Barringer, K. Zorn, J.E. M. Stach, M. P. Crump, M. A. Hayes, M. W. van der Kamp, P. R. Race and C. L. Willis, *Angew. Chem. Int. Ed*., 2023, **62**, e 202213053.

(3) Snider, B. B.; Zou, Y. *Org. Lett.* 2005, **7**, 4939–4941.
